# The phosphoproteomic landscape of the neurological manifestations in tuberous sclerosis complex

**DOI:** 10.64898/2026.03.10.698104

**Authors:** Marie Girodengo, Simeon R. Mihaylov, Katarzyna Klonowska, Laura Mantoan Ritter, Helen R. Flynn, J. Mark Skehel, Elias Bou Farhat, Eleonora Aronica, Matthew White, David J. Kwiatkowski, Sila K. Ultanir, Joseph M. Bateman

## Abstract

Tuberous sclerosis complex (TSC) is a rare disease caused by mutations in *TSC1* and *TSC2,* resulting in activation of mechanistic target of rapamycin complex 1 (mTORC1). Neurological manifestations in TSC patients include epilepsy, autism and intellectual disability. Two types of brain lesions, cortical tubers and subependymal giant cell astrocytomas (SEGAs), cause the majority of neurological manifestations in TSC. We have limited understanding of the molecular changes that occur in tubers and SEGAs and how these contribute to disease pathogenesis. To investigate this, we performed proteomic and phosphoproteomic analysis of TSC patient tuber and SEGA tissue. Tubers showed evidence of alterations in mitochondrial respiration, cytoskeleton organisation and neuronal function. However, we were unable to detect mTORC1 activation in tubers, likely due to the small number of cells with complete inactivation of *TSC1* or *TSC2*. By contrast, SEGAs showed evidence of strong mTORC1 activation and large-scale changes in the proteome and phosphoproteome. SEGAs exhibited increased expression of ribosomal proteins and activation of a neuroinflammatory response. Phosphoproteomics identified 6060 phosphosites within 2154 proteins increased in SEGAs. Phosphorylation of multiple proteins involved in RNA-metabolism, including mRNA splicing, were increased in SEGAs. Consistent with this, we found evidence of extensive alterations in mRNA transcript splicing in SEGA tissue. These data greatly expand the repertoire of known mTORC1 target proteins in the human brain and reveal large-scale mis-regulation of mRNA splicing in SEGAs in TSC.

## Introduction

Tuberous sclerosis complex (TSC) is a dominant genetic disorder characterised by benign tumours in multiple organs resulting from inactivating mutations in either *TSC1* or *TSC2* [1]. The estimated incidence of TSC is between 1 in 6000 and 1 in 13,000 live births worldwide [2–5]. *TSC1* encodes the protein hamartin, which forms a complex with the product of *TSC2*, the GTPase activating protein (GAP) tuberin. Multiple organs can be affected in TSC including benign growths, or hamartomas, in the kidneys (angiomyolipomas and renal cysts), heart (cardiac rhabdomyomas), skin (e.g. angiofibromas) and the lungs (lymphangioleiomyomatosis) [6–8]. Neurological manifestations, which have the greatest morbidity, occur in more than 90% of TSC patients, including infantile spasms, intractable epilepsy, autism and cognitive disability, which often begin in infancy [9, 10].

TSC patients develop three main types of neurological malformations: cortical tubers, subependymal nodules (SENs) and subependymal giant cell astrocytomas (SEGAs). Foetal ultrasound and MRI have shown that cortical tubers and SENs develop during gestation weeks 10 to 20 [11, 12]. Tubers, which typically occur in the cerebral cortex, are present in 90% of TSC patients [10, 13, 14]. Tubers with associated refractory epilepsy are targeted for surgical resection, which reduces seizure burden in most cases [15]. Histologically, tubers contain giant cells, thought to derive from neuroepithelial progenitor cells that sustain a second hit in TSC1/TSC2, and neurons with abnormal morphology and identity compared to the surrounding cortical layer [16, 17]. SENs form in the region surrounding the ventricles, are present in around 85% of TSC patients and arise during foetal development and in the neonatal period [15]. SEGAs are derived from SENs that have proliferative capacity, and can cause hydrocephalus, increased intracranial pressure, seizures, and death [18]. SEGAs typically develop during childhood [13, 19]. Giant cells thought to derive from neuroepithelial progenitor cells are a hallmark of SEGAs, which contain few NeuN positive cells and express astrocyte-like and neural stem cell genes [20].

Seizures in TSC are often refractory to treatment with anti-epileptic drugs and are a significant factor in cognitive outcome. Surgical resection is often the only effective treatment for these patients. Neurodevelopmental and psychiatric disorders including developmental delay, intellectual disability and ASD, neuropsychological deficits, school and occupational difficulties, known as TSC-Associated Neuropsychiatric Disorders (TAND), are also common in TSC [21]. Detailed understanding of the molecular pathogenesis of TSC brain lesions may lead to novel therapeutic strategies.

Nearly all TSC patients have a detectable loss-of-function germline heterozygous or mosaic mutation in *TSC1* or *TSC2* [22], with loss of heterozygosity (LOH) readily identifiable in SEN/SEGA tissue but not in tubers [23–30]. Whole exome sequencing identified second hit somatic mutations in 65% of SEN/SEGAs and 35% of cortical tubers from TSC patients [31]. Laser microdissection of giant cells from TSC patient tuber tissue resulted in consistent detection of somatic mutations that were not detectable in whole tuber sections [32].

TSC1/2 along with TBC1D7 form the TSC protein complex that regulates the mechanistic target of rapamycin (mTOR) pathway. mTOR is a 288 kDa serine/threonine kinase that forms two complexes (mTORC1 and mTORC2) that act as key regulators of processes including nutrient sensing, growth control, autophagy, cytoskeletal dynamics and lipogenesis [33, 34]. Bi-allelic inactivating mutations in TSC1 or TSC2 cause loss of the GAP activity of TSC2 towards the small GTPase Rheb resulting in hyperactivation of mTORC1 signalling [33]. Although a handful of mTORC1 substrates are well characterised, and a recent analysis of the literature identified 57 direct mTORC1 substrates [35], phospho-proteomic studies in mammalian cell lines identified 85-174 proteins whose phosphorylation was regulated by mTOR [36, 37]. Much less is known about mTORC1 targets in the brain, although phosphorylation of eukaryotic translation initiation factor 4E (eIF4E)-binding protein 1/2 (4E-BP1/2) and ribosomal protein S6 (RPS6) are known to be regulated by mTORC1 activity in the developing murine nervous system [38, 39]. Originally identified as an mTORC1 pathway component in the *Drosophila* nervous system [40, 41], the zinc finger/RING domain protein Unkempt is an mTORC1 substrate that physically interacts with Raptor and regulates cognitive flexibilty in mice [42, 43]. However, the targets of mTORC1 in the brain of TSC patients have not been systematically identified.

Rapamycin and derivatives, including everolimus, are allosteric inhibitors of mTORC1 that effectively block its kinase activity for many but not all substrates. Rapamycin and/or everolimus have been shown to be effective for all of the tumour manifestations seen in TSC, including kidney angiomyolipoma, cardiac rhabdomyoma, facial angiofibroma, and lymphangioleiomyomatosis [44, 45]. In a single arm clinical trial, everolimus treatment led to SEGA response, defined as a ≥50% reduction in the sum ot the volume of SEGA lesions, in 58% of 111 patients with TSC and SEGA, that was durable for over four years [46]. Other patients on this trial had smaller degrees of SEGA volume reduction, and SEGA progression on therapy occurred in only about 10% of subjects.

Here we used unbiased tandem mass tag (TMT) labelling and quantitative proteomics and phosphoproteomics to interrogate TSC patient surgical tuber and SEGA tissue. We found significant changes to the proteome in tuber tissue indicating altered mitochondrial processes. Proteomics showed strong evidence of increased ribosome biogenesis and activation of neuroinflammatory processes in SEGA tissue. Changes in the phosphoproteme in tubers indicated perturbed cytoskeleton organisation and neuronal function, but we did not detect evidence of mTORC1 activation in tubers. By contrast, phosphorylation of canonical targets of mTORC1 was strongly increased in SEGA tissue and we identified over 2000 novel mTORC1 targets whose phosphorylation was significantly increased in SEGAs. Gene ontology analysis indicated concerted increases in phosphorylation of RNA-metabolism/mRNA splicing proteins in SEGAs. Moreover, analysis of RNA-sequencing data showed large-scale changes in mRNA transcript splicing in SEGA tissue, including the transcripts for proteins themselves involved in splicing. Thus, mTORC1 activation results in profound changes to the phosphoproteome in SEGAs and perturbs mechanisms regulating RNA metabolism and mRNA splicing at multiple levels.

## Results

### TSC patient clinical and genetic characteristics

We obtained fresh frozen surgical resection frontal/posterior parietal cortical or temporal lobe tuber tissue from three male and three female TSC patients ranging from 7-20 years of age with a median age of 9.5 years old (Figure 1A, Tables 1 and 6). All TSC tuber patients had epilepsy and three also had autism (Table 1). Genetic analysis showed that five tuber patients had germline pathogenic *TSC2* variants and one patient had a *TSC2* variant of unknown significance (Table 2). The identified germline variants included three small deletions of 4-18 bp, two missense variants, and one large deletion encompassing the entire genomic extent of *TSC2* (exons 1 - 41).

**Figure 1.**
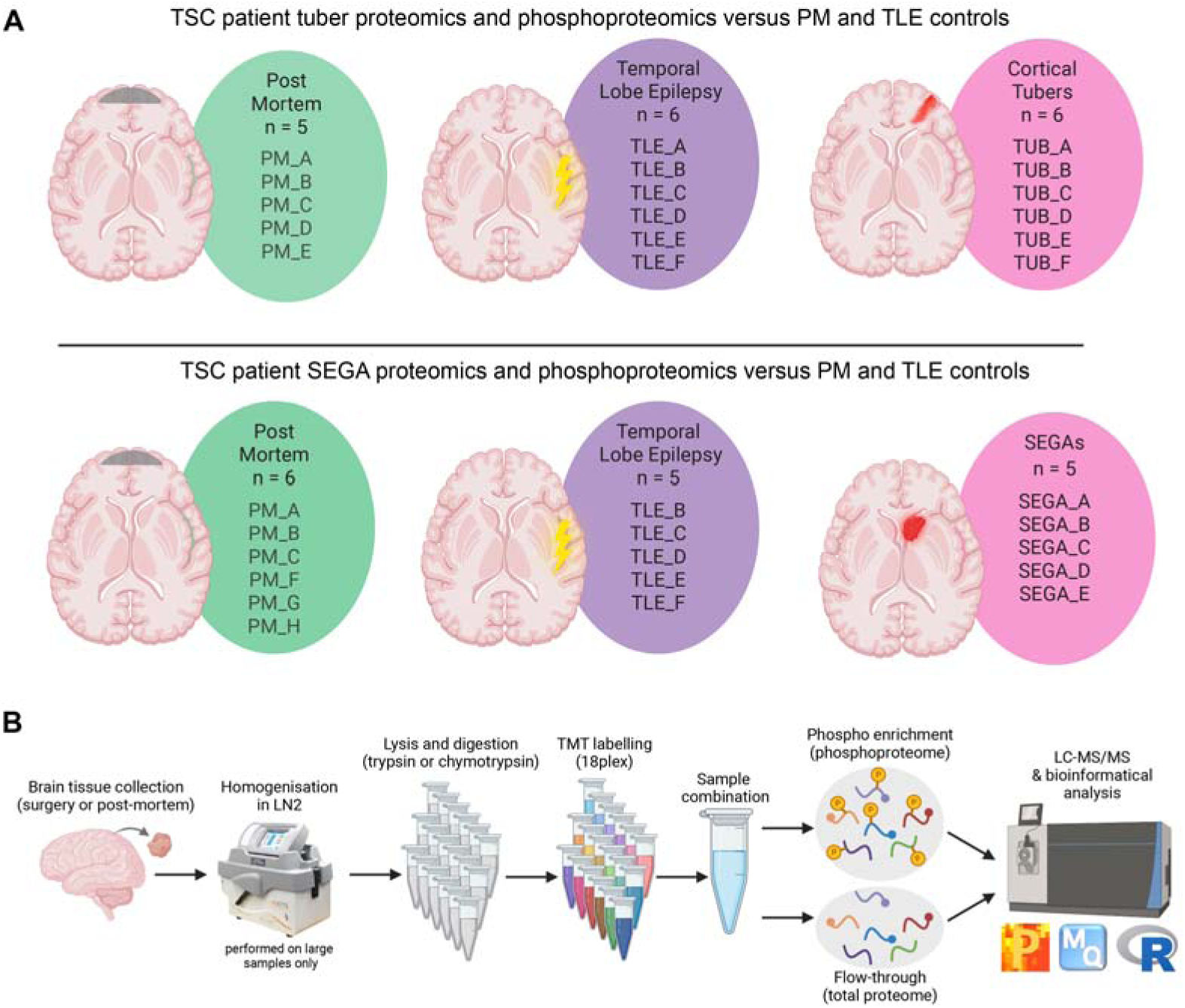
Overview of the experimental conditions and pipeline. (A) Schematic of the TSC patient tissue and control sample sizes for tubers (upper panel) and SEGAs (lower panel). (B) Schematic of the proteomics and phosphoproteomics methodology and analysis pipeline. PM: postmortem controls, TLE: temporal lobe epilepsy controls. PM control samples were age-match to either tuber or SEGA tissue.

**Table 1.**
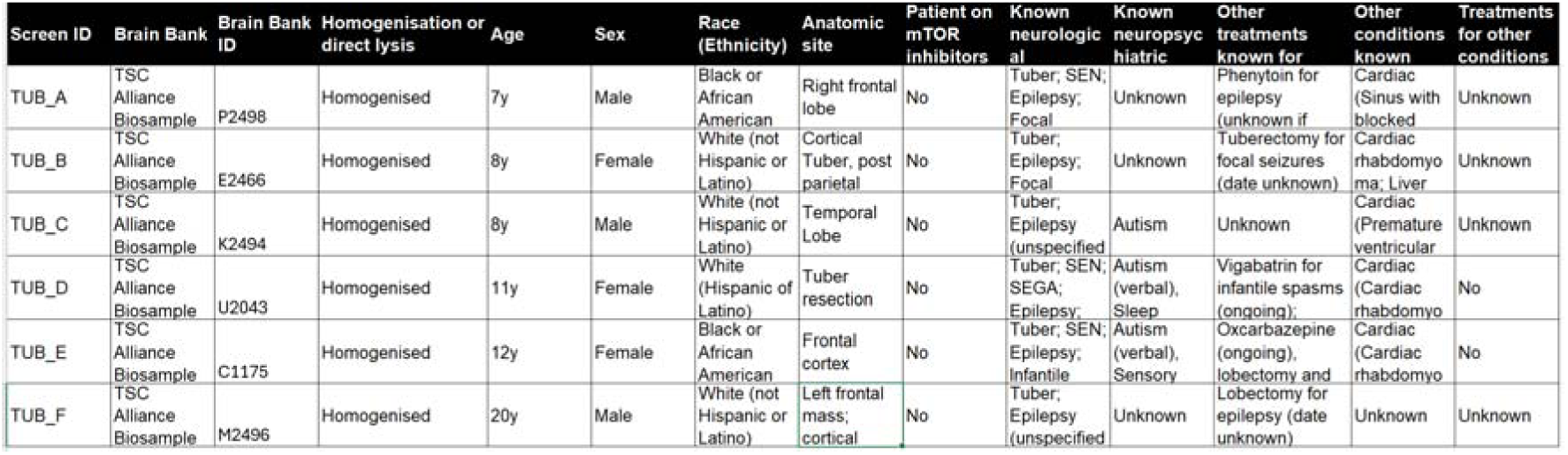
Clinical information on tuber specimen TSC patients.

**Table 2.**
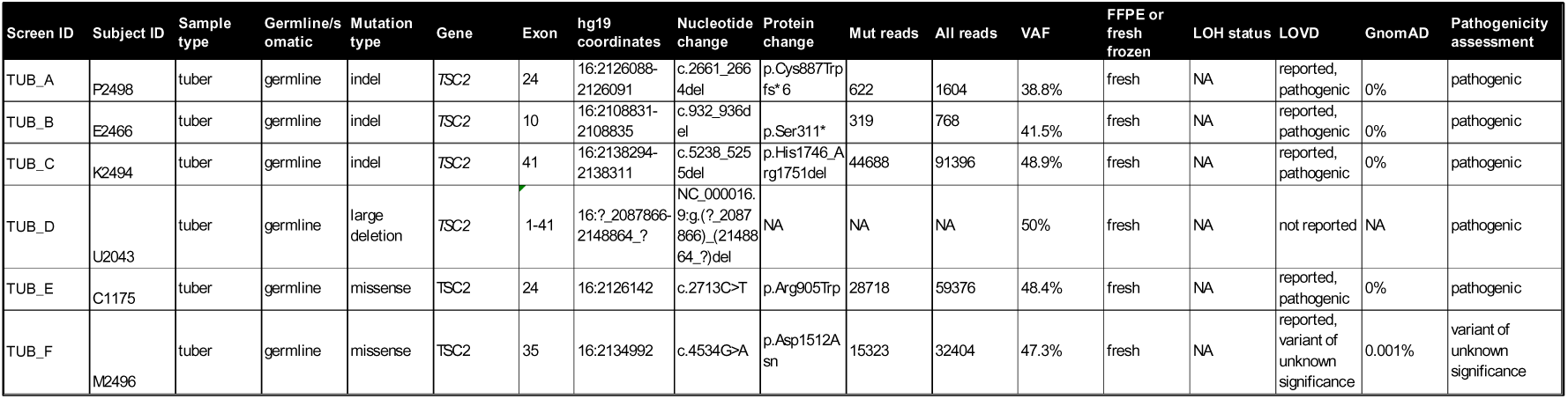
Genetic information on tuber specimen TSC patients.

We obtained fresh frozen surgical resection SEGA tissue from one male and four female TSC patients ranging from 11-25 years of age with a median age of 18 years old (Figure 1A, Tables 3 and 6). Three of the TSC SEGA patients were reported to have epilepsy (Table 3). Genetic analysis was performed for four of the SEGA patients and revealed that all had germline pathogenic *TSC2* variants (Table 4). The *TSC2* variants included two small (1 bp) deletions, one small (2 bp) duplication, and one missense variant. Loss of heterozygosity (LOH) for *TSC2* was identified in 2 of 4 SEGAs (50%) (Table 4).

**Table 3.**
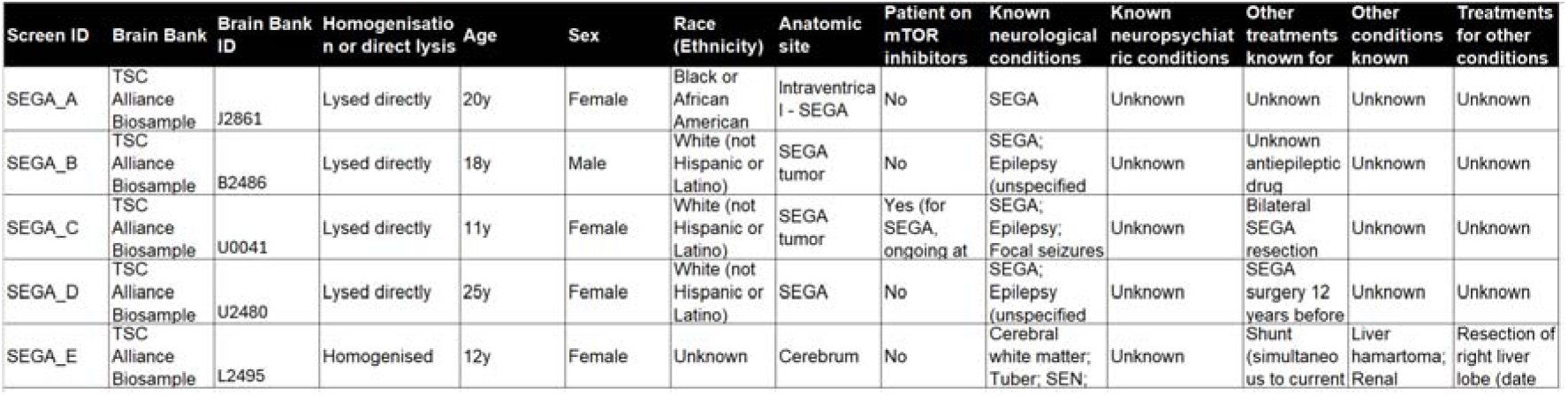
Clinical information on SEGA specimen TSC patients.

**Table 4.** Genetic information on SEGA specimen TSC patients.

| New Screen ID | Subject ID | Sample type | Germline/somatic | Mutation type | Gene | Exon | hg19 coordinates | Nucleotide change | Protein change | Mut reads | All reads | VAF | FFPE or fresh frozen | LOH status | LOVD | GnomAD | Pathogenicity assessment |
| --- | --- | --- | --- | --- | --- | --- | --- | --- | --- | --- | --- | --- | --- | --- | --- | --- | --- |
| SEGA_A | J2861 | SEGA | NA | NA | NA | NA | NA | NA | NA | NA | NA | NA | fresh | NA | NA | NA | NA |
| SEGA_B | B2486 | SEGA | germline | indel | TSC2 | 29 | 16:2129568 | c.3298del | p.Val1100Cysfs*3 | 282 | 653 | 43% | fresh | NO | reported, pathogenic | 0 | pathogenic |
| SEGA_C | U0041 | SEGA | germline | missense | TSC2 | 24 | 16:2126119 | c.2690T>C | p.Phe897Ser | 18803 | 40616 | 46% | fresh | NO | reported, pathogenic | 0 | pathogenic |
| SEGA_D | U2480 | SEGA | germline | indel | TSC2 | 24 | 16:2126078-2126079 | c.2649_2650dup | p.Tyr884Cysfs*11 | 1157 | 1644 | 70% | fresh | YES, TSC2 LOH | not reported | 0 | pathogenic |
| SEGA_E | L2495 | SEGA | germline | indel | TSC2 | 38 | 16:2136792 | c.4910del | p.Lys1637Argfs*35 | 31975 | 44353 | 72% | fresh | YES, TSC2 LOH | not reported | 0 | pathogenic |

As controls we used fresh frozen temporal lobe surgical resection tissue from two male and four female temporal lobe epilepsy (TLE) patients ranging from 29-58 years of age, with a median age of 41.5 years old (Figure 1A, Tables 5 and 6). In addition, as aged-matched controls we used Brodmann area 10 postmortem tissue (PM) cortical tissue from five male and three females who died of non-neurological causes aged between 3-27 years, with a median age of 13 years old (Figure 1A, Tables 5 and 6).

**Table 5.**
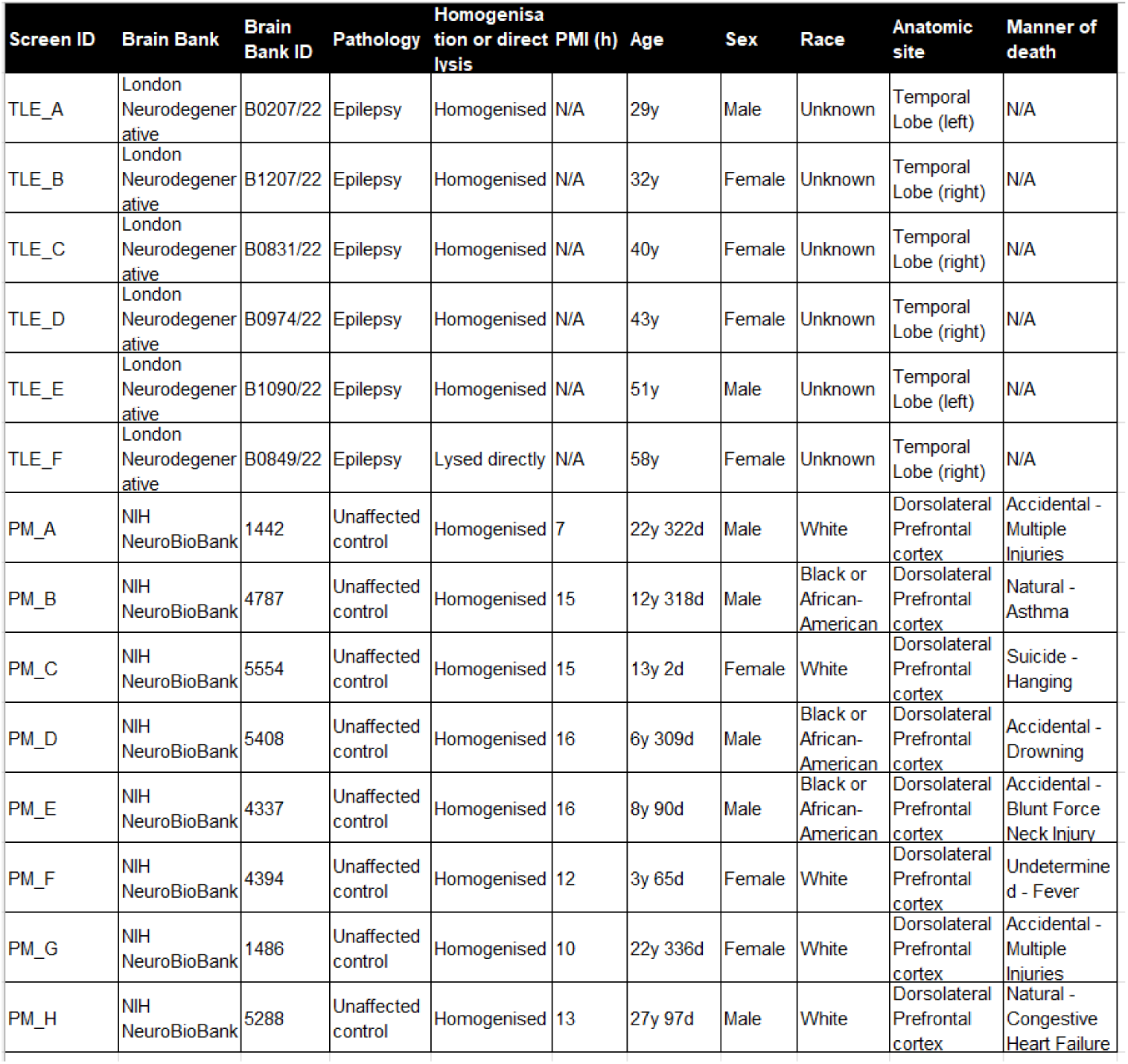
Clinical information on PM and TLE controls.

| Screen ID | Brain Bank | Brain Bank ID | Pathology | Homogenisation or direct lysis | PMI (h) | Age | Sex | Race | Anatomic site | Manner of death |
| --- | --- | --- | --- | --- | --- | --- | --- | --- | --- | --- |
| TLE_A | London Neurodegenerative | B0207/22 | Epilepsy | Homogenised | N/A | 29y | Male | Unknown | Temporal Lobe (left) | N/A |
| TLE_B | London Neurodegenerative | B1207/22 | Epilepsy | Homogenised | N/A | 32y | Female | Unknown | Temporal Lobe (right) | N/A |
| TLE_C | London Neurodegenerative | B0831/22 | Epilepsy | Homogenised | N/A | 40y | Female | Unknown | Temporal Lobe (right) | N/A |
| TLE_D | London Neurodegenerative | B0974/22 | Epilepsy | Homogenised | N/A | 43y | Female | Unknown | Temporal Lobe (right) | N/A |
| TLE_E | London Neurodegenerative | B1090/22 | Epilepsy | Homogenised | N/A | 51y | Male | Unknown | Temporal Lobe (left) | N/A |
| TLE_F | London Neurodegenerative | B0849/22 | Epilepsy | Lysed directly | N/A | 58y | Female | Unknown | Temporal Lobe (right) | N/A |
| PM_A | NIH NeuroBioBank | 1442 | Unaffected control | Homogenised | 7 | 22y 322d | Male | White | Dorsolateral Prefrontal cortex | Accidental - Multiple Injuries |
| PM_B | NIH NeuroBioBank | 4787 | Unaffected control | Homogenised | 15 | 12y 318d | Male | Black or African-American | Dorsolateral Prefrontal cortex | Natural - Asthma |
| PM_C | NIH NeuroBioBank | 5554 | Unaffected control | Homogenised | 15 | 13y 2d | Female | White | Dorsolateral Prefrontal cortex | Suicide - Hanging |
| PM_D | NIH NeuroBioBank | 5408 | Unaffected control | Homogenised | 16 | 6y 309d | Male | Black or African-American | Dorsolateral Prefrontal cortex | Accidental - Drowning |
| PM_E | NIH NeuroBioBank | 4337 | Unaffected control | Homogenised | 16 | 8y 90d | Male | Black or African-American | Dorsolateral Prefrontal cortex | Accidental - Blunt Force Neck Injury |
| PM_F | NIH NeuroBioBank | 4394 | Unaffected control | Homogenised | 12 | 3y 65d | Female | White | Dorsolateral Prefrontal cortex | Undetermined - Fever |
| PM_G | NIH NeuroBioBank | 1486 | Unaffected control | Homogenised | 10 | 22y 336d | Female | White | Dorsolateral Prefrontal cortex | Accidental - Multiple Injuries |
| PM_H | NIH NeuroBioBank | 5288 | Unaffected control | Homogenised | 13 | 27y 97d | Male | White | Dorsolateral Prefrontal cortex | Natural - Congestive Heart Failure |

**Table 6.** Comparison of ages of tuber patients, SEGA patients and controls.

| **Tuber proteomics/phosphoproteomics** |  |  |  |
| --- | --- | --- | --- |
|  | **PM** | **TLE** | **TUB** |
| Number of samples | 5 | 6 | 6 |
| Median age | 13 | 41.5 | 9.5 |
| p value compared to TSC patients | 0.61 | 8.55E-05 |  |
| **SEGA proteomics/phosphoproteomics** |  |  |  |
|  | **PM** | **TLE** | **SEGA** |
| Number of samples | 6 | 5 | 5 |
| Median age | 18 | 43 | 18 |
| p value compared to TSC patients | 0.97 | 7.08E-04 |  |

### Proteomics indicates mitochondrial respiration processes are decreased in tubers

To examine the proteomic and phosphoproteomic landscape in tuber tissue, tuber lysates were digested with trypsin, reduced and alkylated followed by TMT labelling. Labelled samples were pooled together and subjected to phosphopeptide-enrichment using sequential metal oxide affinity chromatography (SMOAC) prior to liquid chromatography tandem mass spectrometry (Figure 1B) [47, 48]. Flow-through material from SMOAC was used for the generation of proteome data (Figure 1B). Proteomic analysis detected 6,058 proteins while phosphoproteomics detected 17,475 phosphopeptides (Figure 2A). Principle component analysis (PCA) of proteins and phosphopeptides showed that four of the tuber samples were well separated from the TLE and PM controls, while two tuber samples (A and B) were separated from the other tuber samples, and clustered with the controls by proteomic analysis (Supplemental figure 1A) and the TLE controls by phosphoproteomic analysis (Supplemental figure 1B). Using a statistical cutoff of p<0.005, the expression level of 1,073 proteins were significantly different between tuber and PM tissue, while 850 proteins were significantly different between tuber and TLE tissue (Figure 2A-C; Supplemental Table S1). The expression of 307 proteins were significantly different between tuber tissue and both PM and TLE tissue (Figure 2A).

**Figure 2.**
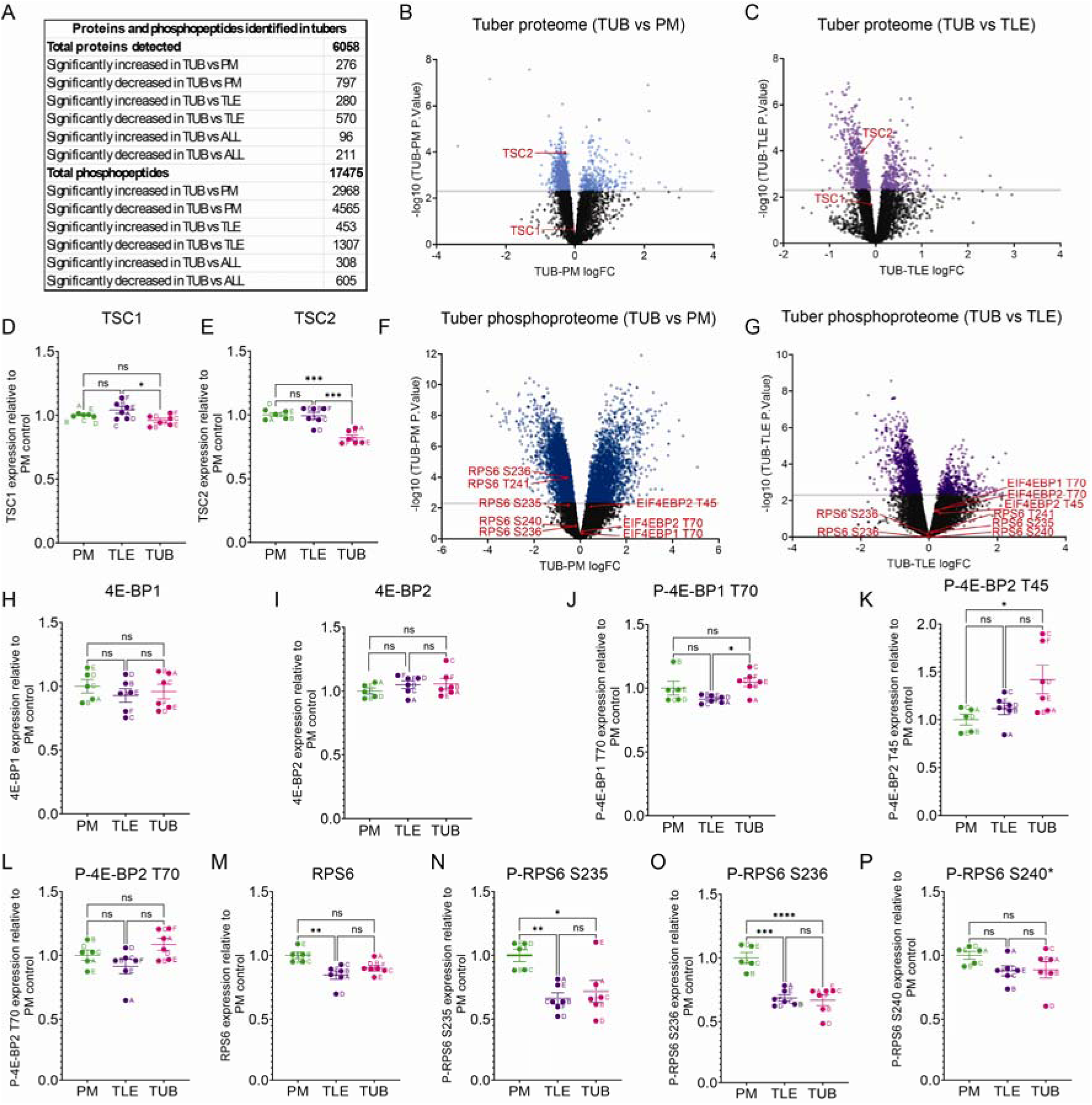
Proteomics and phosphoproteomics of tubers detects reduced TSC1/2 expression but no mTORC1 activation. (A) Summary of the number of proteins and phosphopeptides detected and the number of significantly different proteins and phosphopeptides. (B, C) Volcano plots of TUB versus PM control (B) and TLE control (C) proteomic data. Blue and purple dots represent proteins with significantly altered levels in tubers versus controls. TSC1 and TSC2 are labelled in red. (D, E) TSC1 (D) and TSC2 (E) protein expression levels are slightly reduced in tubers. (F, G) Volcano plots of TUB versus PM control (F) and TLE control (G) phosphoproteomic data. Blue and purple dots represent phosphopeptides with significantly altered levels in tubers versus controls. 4E-BP1/2 (EIF4EBP1/2) and RPS6 phosphopeptides are labelled in red. (H, I) 4E-BP1 (H) and 4E-BP2 (I) protein expression levels are unchanged in tubers. (J-L) Phospho-4E-BP1 T70 and phospho-4E-BP2 T45 and T70 peptide expression levels are unchanged or slightly increased in tubers. (M) RPS6 protein expression levels are unchanged in tubers. (N-P) Phospho-RPS6 S235, S236 and S240 peptide expression levels are unchanged or decreased in tubers. n=5 for PM, n=6 for TLE, n=6 for TUB. Data are represented as mean ± SEM and were analysed using one-way ANOVA. ns not significant, * p < 0.05, **p < 0.01, ***p < 0.001****p < 0.0001.

The proteomic data showed that there was a small but significant decrease in TSC1 expression (6%) between TLE and tuber tissue and a greater decrease in TSC2 expression (17-18%) between tuber and both PM and TLE controls (Figure 2D, E). We also used the proteomic ruler method, which uses the mass spectrometry signal from histones that are proportional to the amount of DNA in the sample, to quantify absolute TSC1 and TSC2 copy number per cell [49]. The protein ruler method showed that TSC1 and TSC2 had similar copy numbers and that the copy number of both proteins was significantly decreased in tubers compared to TLE tissue (23% reduction in TSC1 copy number in tubers versus TLE and 34% reduction in TSC2 copy number in tubers versus TLE; Supplemental Figure S1C, D).

To understand the differences in the protein expression profile of tuber tissue we performed gene ontology (GO) analysis using STRING [50, 51]. From the proteomic data, proteins with significantly increased expression in tuber tissue showed enrichment for the biological processes GO term ‘response to cytokine’, but no molecular function or cellular component terms were significantly enriched (Supplemental Figure S2A). Proteins with significantly decreased expression were dominated by mitochondria-related GO terms including oxidative phosphorylation, aerobic respiration and electron transport chain (Supplemental Figure S2B-D).

### Phosphoproteomics indicates changes in cytoskeleton organisation and neuronal function in tuber tissue

Phosphoproteomic analysis of tuber tissue showed that the level of 7,533 phosphopeptides were significantly different between tuber and PM tissue, while 1,760 phosphopeptides were significantly different between tuber and TLE tissue (Figure 2A, F, G; Supplemental Table S2). 912 phosphopeptides were significantly different between tuber tissue and both PM and TLE tissue (Figure 2A). Proteins with significantly increased phosphorylation in tuber tissue showed enrichment for GO terms involving cytoskeleton organisation and actin filament-based process (Supplemental Figure S2E-G). Proteins with significantly decreased phosphorylation showed enrichment for GO terms involving neuronal functions including neurotransmitter transport, synaptic vesicle cycle and synaptic vesicle membrane (Supplemental Figure S2H, I). These data suggest that cytoskeleton organisation processes are increased, while oxidative metabolism and neuronal processes are reduced in tuber tissue.

### Phosphoproteomics shows no mTORC1 activation in tuber tissue

It was surprising that the GO analysis of the tuber tissue proteomics or phosphoproteomic data did not identify processes or functions relating to the mTOR pathway. Therefore, we next focused on RPS6 and 4E-BP, established targets of mTORC1, whose phosphorylation should be increased in cells that lack TSC1 or TSC2 as a result of activated mTORC1 signalling. Total protein expression levels of 4E-BP1 and 4E-BP2 were unchanged in tubers compared to controls (Figure 2H, I). Neither 4E-BP1 nor 4E-BP2 phosphorylation were significantly increased in the overall dataset (Figure 2F, G). When compared directly, there was a small but significant increase in 4E-BP1 T70 phosphorylation in tubers compared to TLE tissue and in 4E-BP2 T45 phosphorylation compared to PM tissue, but no change in 4E-BP2 T70 phosphorylation (Figure 2J-L). Total protein expression levels of RPS6 were unchanged in tubers compared to controls (Figure 2M). RPS6 phosphorylation was not significantly increased in the overall dataset (Figure 2F, G). When compared directly, RPS6 S235 and S236 phosphorylation were significantly decreased in tuber and TLE tissue compared to PM control tissue and RPS6 S240 phosphorylation was unchanged compared to both controls (Figure 2N-P).

We next compared the tuber phosphoproteomic data with a recently described list of 57 direct mTORC1 substrates consisting of 140 phosphosites (Supplemental Tables S3) [35]. Of the phosphosites detected, none were significantly increased in tuber tissue compared to both PM and TLE controls (Supplemental Tables S3). Thus, consistent with the small decrease in TSC2 expression, using phosphoproteomics we were unable to detect evidence of mTOR pathway activation in TSC patient tuber tissue.

### TSC1 and TSC2 expression are strongly reduced in SEGA tissue

Tubers and SEGAs have distinct anatomical locations and pathology. To understand the proteomic and phosphoproteomic landscape of SEGAs, we used the same experimental pipeline as tuber tissue (Figure 1B), but samples were digested separately with trypsin and chymotrypsin to increase the depth of coverage of peptides lacking the consensus sequence for trypsin. Proteomic analysis using trypsin detected 6,304 proteins (Figure 3A, Supplemental Tables S4, S5). PCA of the trypsin proteomic data showed that the SEGAs were well separated from the PM and TLE controls (Figure 3B).

**Figure 3.**
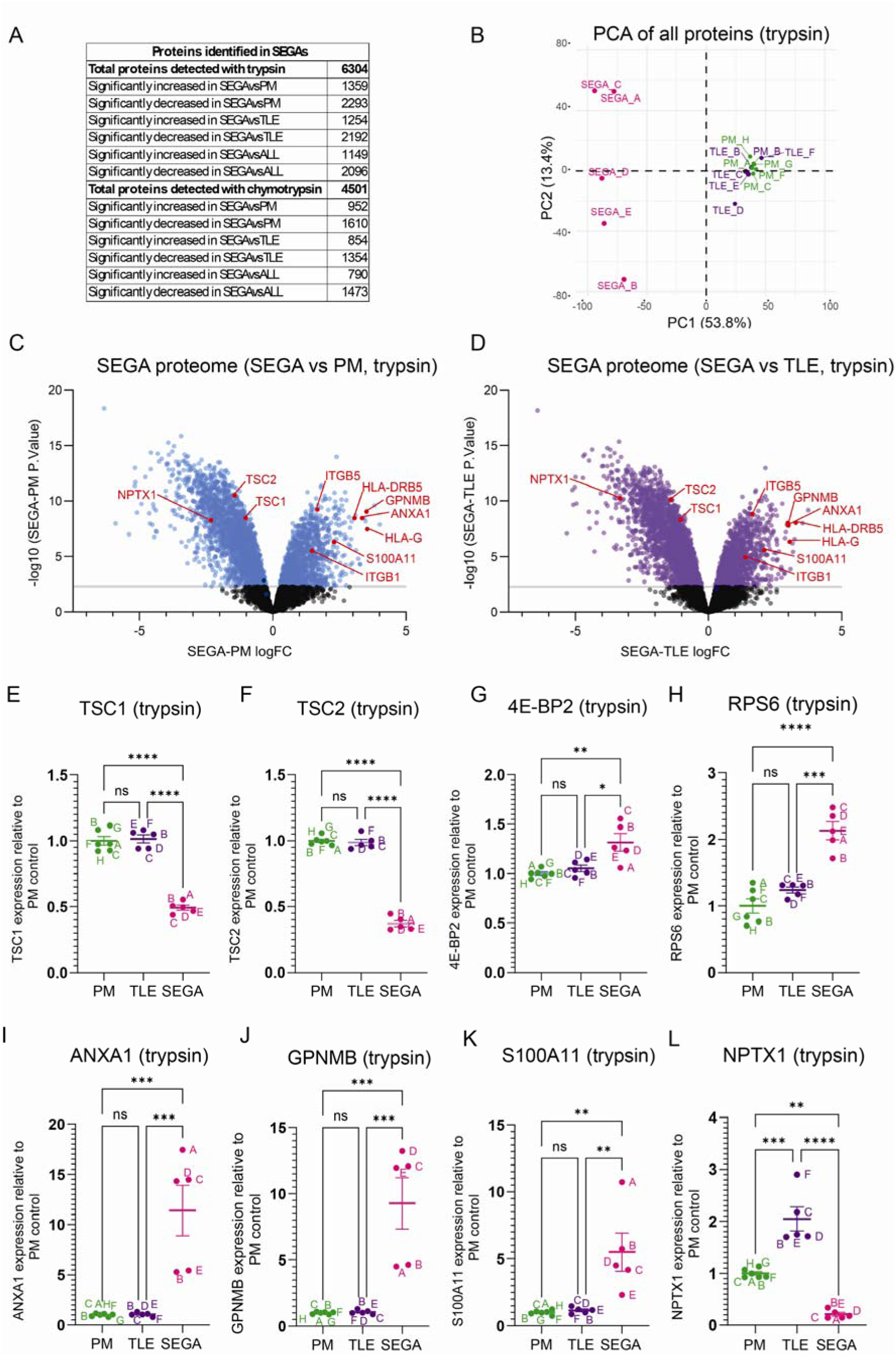
TSC1 and TSC2 expression are strongly reduced in SEGA tissue. (A) Summary of the number of proteins detected and the number of significantly different proteins in the trypsin and chymotrypsin conditions. (B) PCA plot of postmortem (PM), temporal lobe epilepsy (TLE) and SEGA proteomic data in the trypsin condition. (C, D) Volcano plots of SEGA versus PM (C) and TLE (D) trypsin proteomic data. Blue and purple dots represent proteins with significantly altered levels in SEGAs versus controls. TSC1, TSC2, HLA and ITG proteins are labelled in red. (E, F) TSC1 (E) and TSC2 (F) protein expression levels in the trypsin condition are strongly decreased. (G, H) 4E-BP1 (G) and RPS6 (H) protein expression levels in the trypsin condition are increased. (I-K) ANXA1 (I), GPNMB (J) and S100A11 (K) protein expression levels in the trypsin condition are strongly increased. (L) NPTX1 protein expression levels in the trypsin condition is strongly decreased. n=6 for PM, n=5 for TLE, n=5 for SEGA. Data are represented as mean ± SEM and were analysed using one-way ANOVA. ns not significant, * p < 0.05, **p < 0.01, ***p < 0.001****p < 0.0001.

The expression of TSC1 and TSC2 in SEGA tissue were strongly decreased. With trypsin, TSC1 expression was decreased by 45% and TSC2 expression was decreased by 55% in SEGAs compared to PM controls (Figure 3C-F). The protein ruler method also showed a strong decrease in TSC1 and TSC2 protein copy number in SEGAs compared to PM and TLE controls (Supplemental Figure S1E, F). Moreover, the total protein levels of mTORC1 targets 4E-BP2 and RPS6 were significantly increased by 30% and 110% respectively, compared to the PM control, in SEGA tissue (Figure 3G, H).

Proteomic analysis with chymotrypsin showed similar results to trypsin. With chymotrypsin we detected 4,501 proteins, which were well separated from PM and TLE controls by PCA (Figure 3A, Supplemental figure S3A). TSC1 expression was decreased by 30% and TSC2 expression was decreased by 84% in SEGAs compared to PM controls with chymotrypsin (Supplemental figure S3B, C).

Transcriptomic analysis previously identified 50 genes with significantly altered expression in SEGA tissue including ANXA1, GPNMB, and S100A11 and these proteins were also shown to be increased in SEGAs by immunostaining [52]. In our total proteome analysis, ANXA1, GPNMB, and S100A11 were three of the most significantly increased proteins in SEGA tissue (Figure 3C, D, I-K; Supplemental Table S4). This study also found decreased expression of genes involved in nervous system development, including NPTX1. NPTX1 was one of the proteins with the most significantly decreased expression in our SEGA tissue (Figure 3C, D, L; Supplemental Table S4).

Together, these proteomic data show that SEGAs exhibit changes in protein expression consistent with previous studies and show a much greater decrease in TSC1 and TSC2 expression compared to tuber tissue.

### Proteomic analysis shows SEGA tissue exhibits increased ribosomal biogenesis and an inflammatory response

GO analysis of proteins with significantly increased expression in SEGA tissue compared to both PM and TLE controls, combined from the trypsin and chymotrypsin datasets, showed the greatest enrichment for GO terms around cytoplasmic translation and the ribosome (Figure 4A-C). In fact, 33 large ribosomal subunits and 27 small ribosomal subunits had increased expression in SEGA tissue, including RPL37 and RPS27L (Figure 4D, E; Supplemental Tables S4, S5), consistent with the role of mTORC1 in promoting protein translation. GO biological process analysis also showed enrichment for terms antigen processing and presentation (Figure 4A) and the expression of 12 MHC human leukocyte antigen (HLA) proteins were increased in SEGAs, including HLA-G and HLA-DRB5 that were increased more than 10-fold (Figure 3D, E; Figure 4F-H, Supplemental Tables S4, S5). These data indicate that, in addition to promoting protein synthesis, SEGA tissue exhibits a neuroinflammatory response.

**Figure 4.**
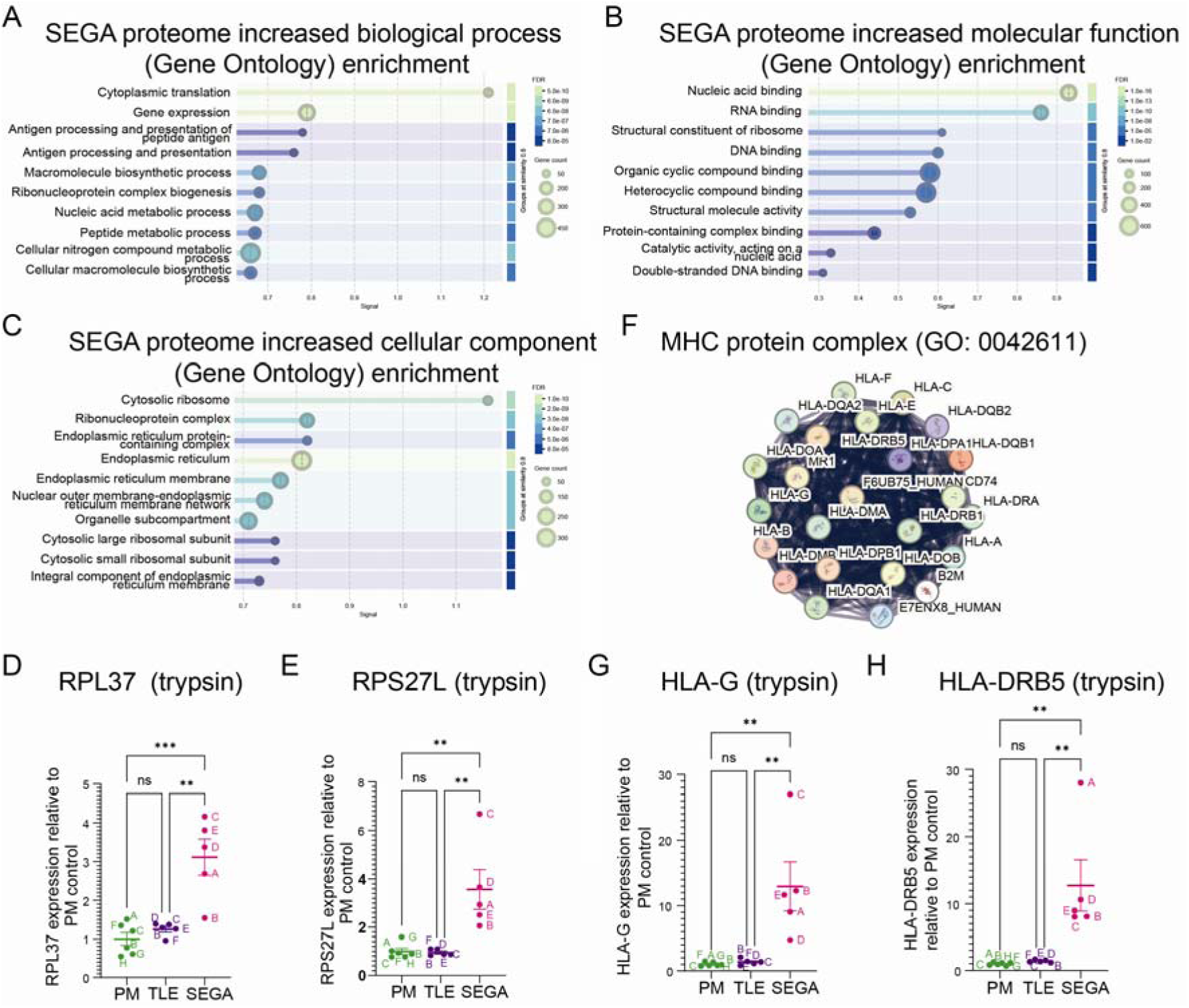
Proteomics of SEGA tissue identities increased ribosomal protein expression and a neuroinflammatory response. (A-C) GO analysis of biological process (A), molecular function (B) and cellular component (C) of proteins with significantly increased expression in SEGAs shows strong enrichment for inflammatory processes. (D, E) RPL37 and RPS27L expression in the trypsin condition are increased in SEGAs. (F) Network nodes representation of MHC protein complex (GO: 0042611) proteins with significantly increased expression in SEGAs. (G, H) HLA-G and HLA-DRB5 expression in the trypsin condition are increased in SEGAs. n=6 for PM, n=5 for TLE, n=5 for SEGA. Data are represented as mean ± SEM and were analysed using one-way ANOVA. ns not significant, * p < 0.05, **p < 0.01, ***p < 0.001****p < 0.0001.

Proteins with decreased expression in SEGA tissue were highly enriched for GO classes including chemical synaptic transmission, presynapse, postsynapse and somatodendritic compartment (Supplemental Figure S3D-F). The enrichment of neuron-related GO terms in the proteins with decreased expression potentially reflects the unique cellular composition of SEGA tissue, compared to the neuron-rich temporal lobe TLE and cortical PM tissue controls.

### Phosphoproteomics detects activation of mTORC1 signalling in SEGA tissue

Phosphoproteomic analysis of trypsin and chymotrypsin digested tissue was used to analyse changes in protein phosphorylation in SEGAs. Phosphoproteomic analysis using trypsin detected 24,861 phosphopeptides and 12,248 phosphopeptides with chymotrypsin (Figure 5A; Supplemental Tables S6, S7). PCA of the phosphoproteomic data showed that the SEGA tissue samples were very well separated from the PM and TLE controls (Figure 5B, Supplemental figure S3G).

**Figure 5.**
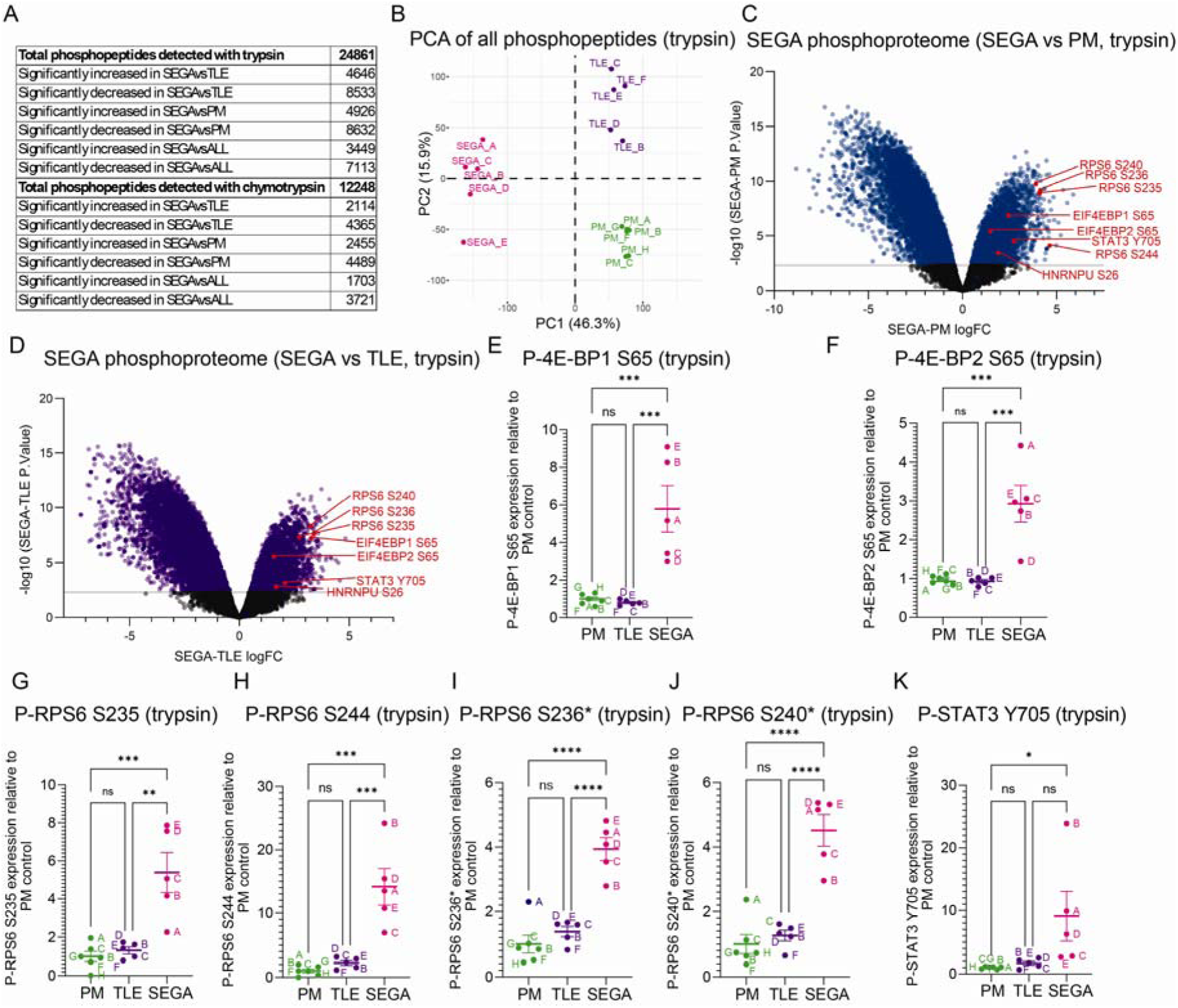
mTORC1 signalling is activated in SEGAs. (A) Summary of the number of phosphopeptides detected and the number of significantly different proteins in the trypsin and chymotrypsin experiments. (B) PCA plot of postmortem (PM), temporal lobe epilepsy (TLE) and tuber (SEGA) phosphoproteomic data in the trypsin condition. (C, D) Volcano plots of SEGA versus PM (C) and TLE (D) trypsin phosphoproteomic data. Blue and purple dots represent phosphopeptides with significantly altered expression in SEGAs versus controls. 4E-BP1/2 (EIF4EBP1/2), RPS6, STAT3 and HNRNPU phosphopeptides are labelled in red. (E, F) Phospho-4E-BP1 S65 (E) and phospho-4E-BP2 S65 (F) phosphopeptide expression levels in the trypsin condition are increased in SEGAs. (G-J) Phospho-RPS6 S235 (G), S244 (H), S236* (I) and S240* (J) phosphopeptide expression levels in the trypsin condition are increased in SEGAs. Asterisks indicate peptides with 1 additional phosphorylated residues. (K) Phospho-STAT3 Y705 phosphopeptide expression levels in the trypsin condition is increased in SEGAs. n=6 for PM, n=5 for TLE, n=5 for SEGA. Data are represented as mean ± SEM and were analysed using one-way ANOVA. ns not significant, * p < 0.05, **p < 0.01, ***p < 0.001****p < 0.0001.

In contrast to tuber tissue, the phosphorylation of established targets of mTORC1 was significantly increased in SEGA tissue compared to controls in the overall datasets (Figure 5C, D). When compared directly, phosphorylation of 4E-BP1 at T37, T46, S65, T70 and 4E-BP2 at S65 were all significantly increased in SEGAs (Figure 5E, F; Supplemental Tables S6, S7). RPS6 phosphorylation was significantly increased at S235 and S244 (Figure 5C, D, G, H; Supplemental Table S6), and also at S236 and S240 but only in peptides that were phosphorylated at one additional serine residue (Figure 5C, D, I, J; Supplemental Table S6), suggesting priming phosphorylation was required for phosphorylation at these residues. Moreover, phosphorylation of STAT3 Y705, which is strongly increased in giant cells of SEGAs [25], was increased in SEGAs on average 9-fold compared to PM controls (Figure 5C, D, K; Supplemental Table S6).

To further validate the SEGA phosphoproteomic data, we combined the trypsin and chymotrypsin phosphoproteomic datasets of significantly increased phosphopeptides compared to controls, and compared this combined SEGA phosphosite dataset with 57 previously reported direct mTORC1 substrates [35]. 40 of 57 (70%) phosphosites, corresponding to 20 proteins, were significantly increased in the SEGA dataset with a median fold change of 3.06 (Supplemental Table S8). Together with the significant increase in P-4E-BP1/2 and P-RPS6 levels, these data provide strong evidence that phosphoproteomics detects activation of mTORC1 signalling in TSC patient SEGA tissue.

We therefore used log FC ≥ 1 (and p<0.005) as a cutoff to identify all potential mTORC1 targets in the SEGA phosphosite dataset. Since some previously reported direct mTORC1 substrate phosphosites were significantly increased compared to either the PM or TLE control but not both (Supplemental Table S8), we included phosphosites that were significantly increased (p<0.005) compared to either control with a FC ≥ 1. Using these criteria, there were 6060 phosphosites within 2154 proteins that were increased in SEGA tissue (p<0.005, logFC ≥1, Supplemental Table S9 and S10). mTOR has a strong bias towards serine as the phosphoacceptor residue (77% serine, 23% threonine), and a strong bias towards proline at the +1 position with glutamic acid, phenylalanine, tyrosine, and glutamine also favoured somewhat at +1 [35, 36]. Analysis of SEGA phosphosites showed that serine was the dominant phosphoacceptor residue (85%), with proline most common (29%) at the +1 position and some enrichment for glutamic acid, leucine, serine and aspartic acid (Figure 6A). These data suggest that many of the proteins whose phosphorylation is increased in SEGAs are direct mTORC1 substrates.

**Figure 6.**
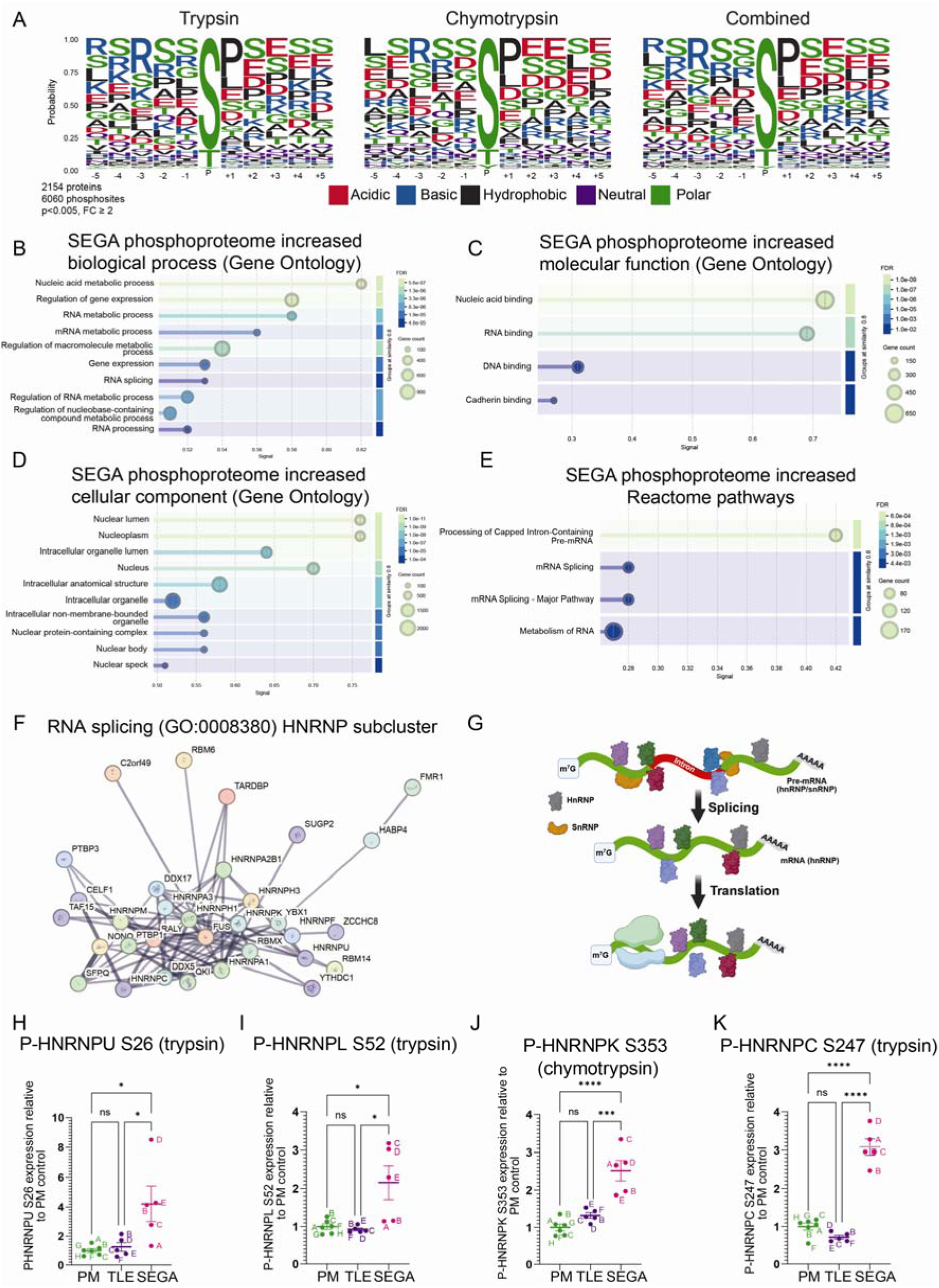
Phosphorylation of proteins regulating RNA metabolic processes are increased in SEGAs. Residue enrichment motif analysis of the significantly increased SEGA phosphopeptide dataset phosphosites in the trypsin condition, chymotrypsin condition and both conditions combined. (B-D) GO analysis of biological process (B), molecular function (C) and cellular component (D) of proteins represented by phosphopeptides with significantly increased expression in SEGAs shows strong enrichment for RNA-related processes. (E) Reactome pathways of proteins represented by phosphopeptides with significantly increased expression in SEGAs. (F) Network nodes representation of RNA splicing (GO:0008380) proteins represented by phosphopeptides with significantly increased expression in SEGAs, HNRNP subcluster. (G) Schematic of the role of hnRNPs in mRNA splicing and translation. (H-K) phospho-HNRNPU S26 (H), (I), phospho-HNRNPL S52 (J), phospho-HNRNPK S353 (K) and phospho-HNRNPC S247 (K) phosphopeptide levels in either the trypsin or chymotrypsin conditions are increased in SEGAs. n=6 for PM, n=5 for TLE, n=5 for SEGA. Data are represented as mean ± SEM and were analysed using one-way ANOVA. ns not significant, * p < 0.05, **p < 0.01, ***p < 0.001****p < 0.0001.

### Phosphorylation of RNA metabolism proteins is increased in SEGAs

GO analysis of the significantly increased SEGA phosphopeptide dataset (Supplemental Table S10) showed robust enrichment for nucleic acid metabolic process/nucleic acid binding biological processes and molecular functions as well as nucleus related cellular components (Figure 6B-D). In particular, RNA metabolic processes, RNA processing and RNA binding GO terms were very highly enriched (Figure 6B-D). Moreover, Reactome pathways analysis showed enrichment for processing of capped intron containing pre-mRNA, metabolism of RNA and mRNA splicing (Figure 6E).

Many phosphopeptides increased in SEGAs were from proteins involved in mRNA splicing, including 132 of the 208 proteins annotated to GO term ‘RNA splicing’ (GO:0008380) (Figure 6F). mRNA splicing is carried out by the spliceosome, a nuclear-localised multi-megadalton ribonucleoprotein complex involving more than 120 proteins that assembles on pre-mRNA substrates and removes non-coding introns to generate mature mRNAs [53]. Phosphorylation of U2-related spliceosome components (PUF60, SF3B1, SFPQ), pre-mRNA processing factor proteins (e.g. PRPF38A, PRPF38B, PRPF40A), SR-related proteins (SRRM1, SRFBP1, SRPK2) and exon junction complex proteins (ACIN1, RBM8A, CASC3) were all increased in SEGAs (Supplemental Table S9, S10). Phosphorylation of ACIN1 was increased at residues S240 and S243, which are directly phosphorylated by mTORC1 [35, 54]. Moreover, the phosphorylation of 16 heterogeneous nuclear ribonucleoproteins (hnRNPs) was increased in SEGAs (Figure 6F, H-K; Supplemental Table S9, S10). HnRNPs are involved in regulating the maturation of newly formed heterogeneous nuclear RNAs (hnRNAs/pre-mRNAs) into messenger mRNAs, stabilising mRNAs during cellular transport and controlling their translation (Figure 6G) [55]. Remarkably, the activity of five hnRNPs whose phosphorylation was increased in SEGA tissue has been shown to be regulated through phosphorylation at the same residues (Figure 5C, D; Figure 6H-K; Table 7). These data strongly indicate that mTORC1 activation in SEGAs causes concerted changes in the regulation of key proteins involved in mRNA splicing.

**Table 7.** HnRNPs with functionally validated phosphosites that have increased expression in SEGAs. ^a^ Identified in trypsin experiment. ^b^ Identified in chymotrypsin experiment.

| **HnRNP** | **Phosphosite increased in SEGAs** | **How phosphosite regulates function** | **References** |
| --- | --- | --- | --- |
| HNRNPA1 | Serine 6<sup>b</sup> | Facilities mRNA splicesite binding and nuclear export | [91, 92] |
| HNRNPC | Serine 247<sup>a</sup> | Facilities mRNA splicesite binding | [93, 94] |
| HNRNPU | Serine 26<sup>a</sup> | Facilities chromatin binding via N-terminal SAP DNA-binding domain | [94] |
| HNRNPL | Serine 52<sup>a</sup> | Regulates splicing | [95] |
| HNRNPK | Serine 353<sup>b</sup> | Regulates cytoplasmic accumulation and mRNA translation | [96, 97] |

### Evidence of widespread changes to splicing in SEGAs

Based on the increased phosphorylation of proteins involved in RNA-metabolism and mRNA splicing we hypothesised that splicing of mRNAs would be affected in SEGA tissue. Unfortunately, the SEGA tissue available to us for this study was depleted by our proteomic analyses, and RNA-sequencing (RNA-seq) could not be performed for these same samples. Therefore to test this hypothesis, we analysed previously published SEGA tissue RNA-seq data from 17, mostly paediatric, TSC patients and periventricular tissue from 8 (postmortem tissue) controls [56].

We analysed global changes in splicing at the level of isoforms, splicing events and exons. The isoform approach, using IsoformSwitchAnalyzeR [57], identified 3,139 significant isoform switches from 2,526 genes, showing that a substantial subset (∼15%) of genes exhibit significant changes in isoform usage in SEGA tissue versus controls (Figure 7A). The rMATS splicing analysis pipeline is an events based approach that analyses differences in specific types of splice event including alternative 3’ or 5’ splice site, mutually exclusive exons, retained introns and skipped exons [58]. rMATS analysis identified a total of 1,572 significant splicing events with a meaningful ΔPSI in SEGA tissue, highlighting both gains and losses in exon inclusion or splice site usage (Figure 7B, C). Finally, at the exon level we identified 3,817 genes within which differential exon usage was detected, a total of 18,183 exons (Figure 7D). Together, these analyses show clear evidence that SEGA tissue experiences widespread alternative splicing dysregulation, with many significant changes in exon inclusion/exclusion and other splice site usage.

**Figure 7.**
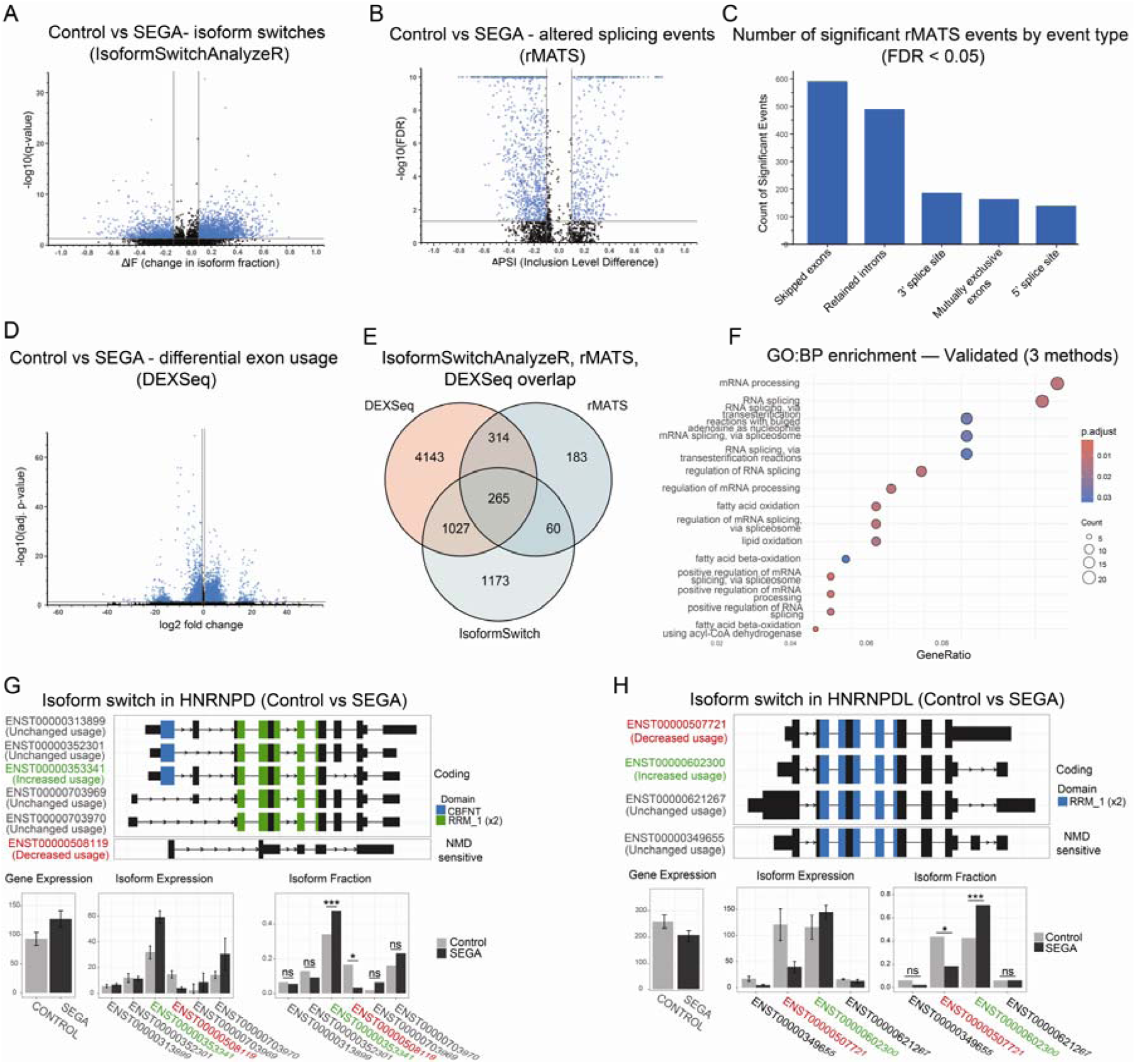
Evidence of large scale alterations in mRNA splicing in SEGAs. (A) Volcano plot showing significantly different isoforms (blue dots) in SEGA tissue versus controls identified using IsoformSwitchAnalyzeR. (B) Volcano plot showing significantly different splicing events (blue dots) identified by rMATS in SEGA tissue versus controls. (C) Number of significant rMATS splicing events of each type in SEGA tissue versus control. (D) Volcano plot showing genes with significantly different exon usage (blue dots) in SEGAs versus controls. (E) Venn diagram showing the number of genes identified using three different methods and genes in common using each method. (F) GO analysis of genes identified using all three methods as having significantly altered splicing in SEGAs versus controls. (G, H) Splicing changes in HNRNPD (G) and HNRNPDL (H) in SEGAs.

We next looked at the overlap between the genes identified using each method of splicing analysis. There was significant overlap between each method, with 1,667 genes identified by at least two approaches (Figure 7E, Supplemental Table S11). Remarkably, biological process GO analysis of genes that had altered splicing in SEGAs validated by all three analysis methods was dominated by enrichment for GO terms around RNA processing and RNA splicing (Figure 7F; Supplemental Table S12). Moreover, splicing of six hnRNPs was altered in SEGA tissue, including isoform switching in HNRNPD and HNRPNPDL (Figure 7G, H; Supplemental Table S11). These data suggest that increased phosphorylation of hnRNPs by mTORC1 activation in SEGAs has global effects on mRNA splicing, including the splicing of hnRNP mRNAs themselves (Figure 8).

**Figure 8.**
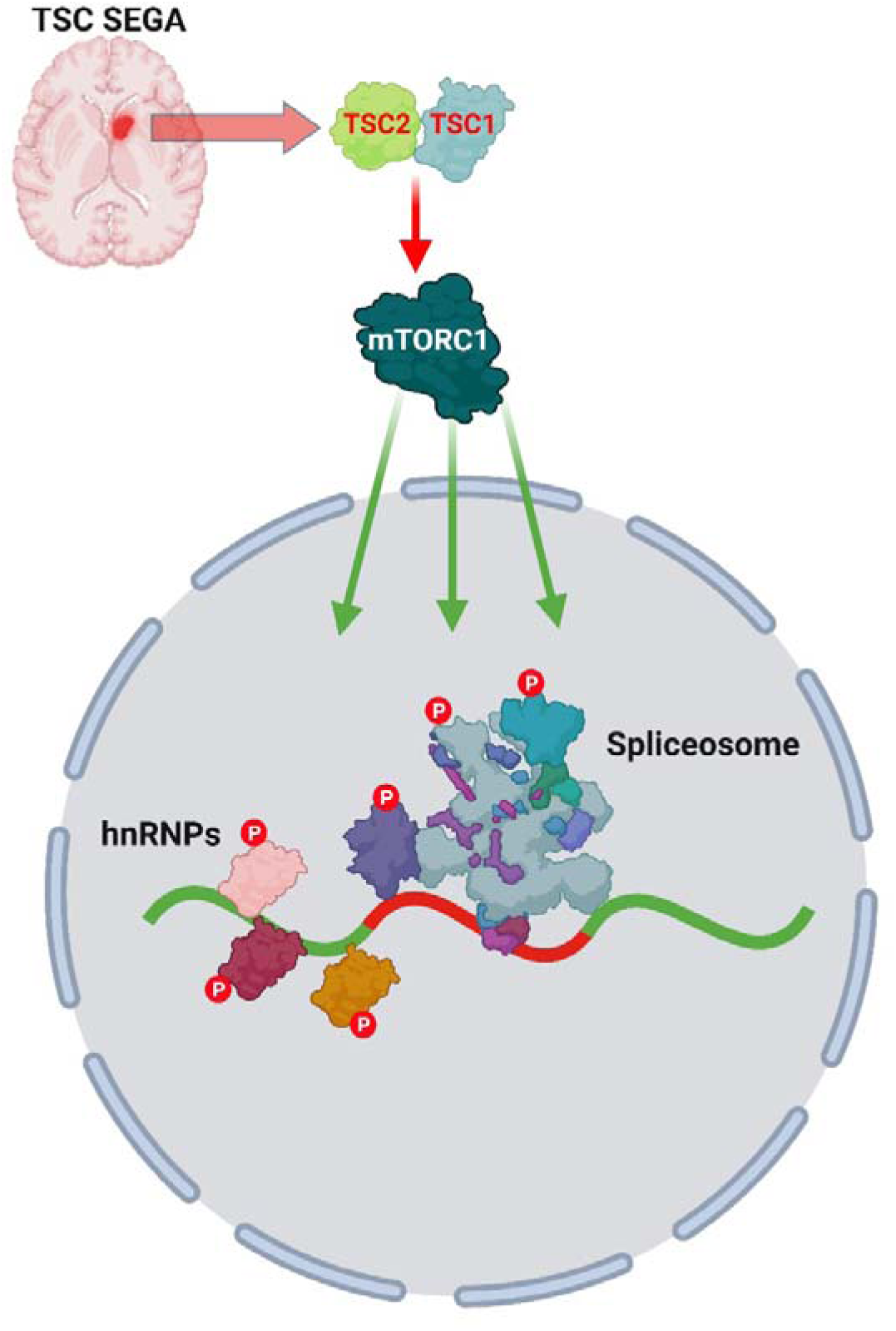
A schematic representing activation of mTORC1 in TSC SEGAs leading to increased phosphorylation of spliceosome components and hnRNPs resulting in widespread alterations in mRNA splicing.

## Discussion

Activation of mTORC1 due to second hit and complete loss of TSC1 or TSC2 function in brain cells during development is the fundamental pathologic process in TSC, resulting in the formation of tubers and SEGAs, a major cause of morbidity and mortality in TSC patients. Our deep molecular phenotyping of TSC patient tuber and SEGA tissue using quantitative proteomics and phosphoproteomics illuminates the complex molecular changes occurring in these lesions. We found evidence for altered cytoskeleton organisation and mitochondrial respiration processes in tubers. SEGA tissue showed much more extensive molecular changes than tubers, with greater decreases in the levels of TSC1 and TSC2 expression and clear evidence of mTORC1 activation. This enabled us to identify a large number of new mTORC1 targets and reveal the phosphorylation-dependent cellular and molecular processes that are disrupted in SEGAs, including RNA metabolism and mRNA splicing. Thus, widespread changes in mRNA splicing may be a significant contributory factor in the neuropathology of SEGAs in TSC. Future functional studies can now be designed to determine the contribution of aberrant mRNA splicing to the neuropathology of TSC.

Previous studies investigating the molecular changes in the brain in TSC have utilised expression profiling or transcriptomics to identify mis-regulated genes and then validated specific candidates at the protein level [24, 31, 52, 56, 59–61]. Our study provides an unbiased snapshot of the levels of over 6,000 proteins and almost 25,000 phosphopeptides from the brain of TSC patients compared to two independent control groups. The choice of control tissue is challenging when analysing TSC patient tissue, particularly with SEGAs as they have a unique cellular makeup. We used postmortem tissue as age-matched controls for the tuber and SEGA tissue. However, protein phosphorylation can be adversely affected by the postmortem interval (PMI) [62, 63]. Although the PMI of the PM controls was relatively short (7-16 hours, Table 5), we used fresh frozen temporal lobe surgical resection tissue as additional controls. This comparison to two different controls greatly enhances the robustness and gives confidence that our data provide a rich resource for deciphering the molecular drivers of the neurological manifestations in TSC.

Tuber tissue had a small decrease in TSC protein expression and we observed relatively few proteins (317) with significantly altered expression in tubers compared to PM and TLE controls. The strongest signal from GO analysis of these 317 proteins was for a decrease in the expression of mitochondrial proteins. TSC2 mutant iPSC-derived neurons have reduced axonal mitochondria, impaired mitochondrial respiration and mitochondrial transport [64]. Moreover, genes with decreased expression in TSC patient iPSC-derived cortical organoids were predominantly enriched for mitochondrial genes and mitochondrial abnormalities were observed using electron microscopy of TSC patient cortical tissue [65]. Compromised mitochondrial function may therefore be a key contributor to the neuropathology of tubers.

The main driver of pathology in TSC patient hamartomas is mTORC1 activation, where enhanced mTOR kinase activity results in increased substrate phosphorylation. Loss of TSC1 and TSC2 causes increased phosphorylation of established mTORC1 targets such as RPS6 and 4E-BP, but there are over 50 known additional direct mTORC1 substrates, and in cultured cells mTORC1 has over 100 putative novel targets [36, 37, 66, 67]. Therefore, fully establishing the molecular landscape of the neurological manifestations in TSC requires omics approaches that directly detect changes in mTORC1 target phosphorylation. The unique genetic makeup of tubers, consisting of a small number of cells with biallelic *TSC1* or *TSC2* gene inactivation [32], likely explain our findings that there is no increase in phosphorylation of established mTORC1 targets. However, the decrease in phosphopeptides from proteins with synaptic functions in tubers is consistent with RNA-seq analysis that found almost all genes with decreased expression in tuber tissue related to synaptic transmission [31]. In future, ultra-low-input spatial tissue proteomics approaches may enable analysis of the proteome and phosphoproteome from a subset of tuber cells with TSC biallelic gene loss [68].

LOH of *TSC1* or *TSC2* has been reported in approximately 80% of SEGAs [23–25, 31] and we were able to confirm that two of the four genetically analysed SEGAs (50%) in our study had *TSC2* LOH. This is consistent with the large decrease in TSC2 expression in SEGAs, indicating that the majority of cells in the tissue have activated mTORC1. SEGAs also had a large number of proteins with significantly altered expression compared to PM and TLE controls (3,245 in SEGAs for the trypsin digest condition compared to 317 in tubers). Moreover, our proteomic analysis showed 5-10 fold increased expression of ANXA1, GPNMB, and S100A11 in SEGAs, while NPTX1 expression was strongly reduced, in agreement with a previous study of SEGA tissue [52]. A potential confound of using SEGA tissue for bulk protein analysis is that changes in the proteome may reflect differences in cellular composition, rather than differences in expression of mTORC1-dependent targets. However, Tyburczy et al., showed that ANXA1, GPNMB, S100A11 expression levels were increased in SEGA tissue and also in primary cell cultures derived from SEGAs, when compared to cultured normal human astrocytes, and these changes were reversed by rapamycin treatment [52]. Although there are differences in cellular composition between the control tissues and SEGAs, these data support the interpretation that the proteomic analysis detected changes in SEGAs driven by mTORC1 activation.

Activation of inflammatory markers has been previously observed in tubers and SEGAs. RNA-seq analysis of SEGAs found that the genes most significantly increased in expression were related to the immune system and inflammation, and activated microglia expressing HLA-DR were observed in SEGA tissue using immunostaining [31]. Moreover, microglial cells expressing HLA-DR cluster around dysplastic neurons in tubers and are abundant in SEGAs [69]. More recently, cells positive for HLA-DP/DQ/DR expression were shown to be significantly increased in SEGAs [70]. Our finding of enrichment for inflammatory proteins in SEGAs, including MHC HLA-G and HLA-DRB5, reinforce the idea that a neuroinflammatory response is activated in SEGAs.

The combined trypsin/chymotrypsin SEGA phosphoproteomic dataset revealed phosphopeptides with increased expression in SEGAs representing 2154 proteins, (Supplemental table S9 and S10), that were distinct from the proteins with increased expression in the SEGA proteomic data, emphasising the importance of understanding the phosphoproteome. Using data from trypsin and chymotrypsin digests significantly increased the depth of the phosphoproteome since around 75% of the phosphosites in the chymotrypsin data were not identified with trypsin. The phosphosites and surrounding residues in the 2154 proteins strongly resembled the consensus from 57 direct mTORC1 substrates, suggesting that many of these proteins are directly phosphorylated by mTORC1 in SEGAs. This pathological mTORC1 phosphoproteome can in future be further interrogated to understand the neurobiology of TSC and identify novel therapeutic targets.

Analysis of the proteins whose phosphorylation was increased in SEGAs showed a dramatic enrichment for factors regulating RNA-metabolism/mRNA splicing. We have previously shown that the RNA binding protein Unkempt is a direct substrate of mTORC1 and phosphorylation modifies the ability of Unkempt to regulate cellular morphogenesis [43]. Unkempt regulates the translation of hundreds of mRNAs and Unkempt phosphorylation is increased in a mouse model of TSC and in SEGA tissue [42, 71–73] (Supplemental table S9 and S10). Aside from Unkempt, mis-regulation of RNA-metabolism/mRNA splicing proteins has not been previously observed in TSC cell or animal models or in TSC patients. Interestingly, phosphoproteomics of myoblasts differentiated from type II diabetes patients had mis-regulated mTORC1 signalling and strong enrichment for proteins relating to mRNA metabolism and mRNA splicing [74]. Moreover, phosphoproteomics of human insulin stimulated myotubes compared to serum starved control cells showed strong enrichment for mRNA splicing functions, increased phosphorylation of spliceosomal proteins and evidence altered mRNA splicing [75]. Therefore, mRNA-metabolism proteins appear to be a primary target of mTORC1 phosphorylation in both muscle tissue and SEGAs.

Our analysis of previously published RNA-seq data from SEGA tissue and controls showed evidence of largescale alterations in mRNA splicing. Remarkably, GO analysis of the genes whose splicing was altered in SEGAs showed dramatic enrichment for genes involved in RNA processing and RNA splicing. This is not unprecedented, as intrinsically disordered domains in hnRNPs have been shown to contribute to the regulation of alternative splicing and are often themselves regulated through alternative splicing [76, 77]. These data indicate that mTORC1 activation in SEGAs results in mRNA splicing being targeted at two stages: (i) phosphorylation of the proteins that regulates splicing, (ii) changes in the splicing of mRNAs encoding RNA processing and RNA splicing proteins.

In sum, our SEGA phosphoproteome data and splicing analyses suggest that mTORC1 activation results in concerted perturbations to RNA-metabolism and mRNA splicing in SEGAs in the TSC patient brain (Figure 8).

## Methods

### TSC patient and control tissue

TSC patient tissue was from the Tuberous Sclerosis Alliance Biosample Repository. TLE patient tissue from the London Neurodegenerative Diseases brain bank. PM tissue was from the NIH Neurobiobank.

### Tissue preparation and lysis

Most samples were homogenised in liquid nitrogen using a 6875D Dual Freezer/Mill (Spex SamplePrep). Each frozen sample was placed inside a mid-size grinding vial (6881C20, Cole-Parmer) alongside a stainless-steel impactor (6881P, Cole-Parmer). Grinding vials were closed with stainless-steel plugs (6881E, Cole-Parmer). Vials, impactors and plugs were all kept on dry ice for at least 15mins prior to manipulation to ensure samples remained frozen. Closed vials were then transferred to the liquid-nitrogen-filled electromagnetic chamber of the freezer mill. Samples were ground using the following programme, with steps 1-3 repeated 6 times: (1) pre-cooling: 2mins; (2) grinding: 2mins, 15 cycles per second (i.e. 30 impacts per second); (3) cooling: 2mins. Powdered homogenates were then transferred to 50mL Falcon tubes using spatulas. Tubes and spatulas were kept on dry ice for at least 15mins before transfer to ensure samples remained frozen. Vials, impactors, plugs and spatulas were thoroughly rinsed with 70% ethanol and double distilled water (ddH2O) between samples. Half of the resulting powdered tissue was lysed in urea buffer for use in phosphoproteomic screens. Some samples of unusually small sizes were lysed directly to avoid any loss associated with the homogenisation process. All homogenised powdered tissue and urea lysates were kept at −80°C.

Powdered tissue was lysed using urea-based lysis buffer containing 8 M urea, 50 mM HEPES (pH 8.2), 10 mM glycerol-2-phosphate, 50 mM sodium fluoride, 5 mM sodium pyrophosphate, 1 mM EDTA, 1 mM EGTA, 1 mM sodium vanadate, 1 mM DTT, 1X cOmplete protease inhibitor cocktail (Roche), 1X phosphatase inhibitor cocktail 3 (Sigma), and 500 nM okadaic acid. Protein concentrations were quantified using Pierce BCA assay. 150 µg of protein per sample for tubers and controls was used, while 100 µg of protein per sample for SEGAs and controls was utilised for TMTpro multiplexed quantitative proteomics.

### Protein reduction, alkylation, and digestion

Protein reduction was performed with 10 mM DTT at 56°C for one hour, followed by alkylation with 20 mM iodoacetamide (IAA) at room temperature in the dark for 30 minutes. The reaction was quenched with 20 mM DTT and diluted to a final urea concentration of less than 2 M with 50 mM HEPES (pH 8.5). Proteins were digested overnight at 37°C with LysC (4.44 µg per sample; Lysyl endopeptidase, 125-05061, FUJIFILM Wako Chemicals) and Trypsin (11.11 µg per sample; MS grade, 90058, ThermoFisher Scientific). In parallel, the SS samples were processed twice – once with LysC and trypsin digest and once with chymotrypsin digest. For the chymotrypsin digest, the final urea concentration was adjusted to less than 1 M with 50 mM HEPES (pH 8.5) and supplemented with 10 mM CaCl_2_. Each sample was then digested with 5.56 µg chymotrypsin (Sequencing grade, Promega, V1061) shaking at 25°C overnight.

### Sample cleanup and TMT labelling

After digestion, samples were quenched with 1% trifluoroacetic acid (TFA) and cleaned with Nest Group BioPureSPN MACRO (Proto 300 C18; Part# HMM S18V) before being freeze-dried using refrigerated benchtop vacuum concentrator. The samples were then labelled with TMTpro 18plex Isobaric Label Reagent Set (0.5 mg per tag, A44522 + A52048, ThermoFisher Scientific; TS LOT XG350092 + XD345167, SS LOT XH351216 + XE347834) according to the manufacturer’s instructions. Labelling efficiency and mixing accuracy were assessed via LC-MS/MS analysis on an Orbitrap Eclipse Tribrid mass spectrometer using with a 60-minute HCD MS2 fragmentation method. Labelling efficiency exceeded 99% for all samples.

### Phosphopeptide enrichment and fractionation

Labelled samples were quenched with hydroxylamine, pooled, partially vacuum-dried, acidified to pH ∼2.0, and cleaned using C_18_ Sep Pak 1cc Vac, 50 or 100 mg bed volume (Waters). A mixing check was performed via LC-MS/MS analysis with a 240-minute HCD MS2 fragmentation method. Sequential enrichment of Metal Oxide Affinity Chromatography (SMOAC) (phosphopeptide enrichment) was carried out using high-select TiO_2_ (Thermo Scientific, A32993) followed by Fe-NTA (Thermo Scientific, A32992) phospho-enrichment columns following manufacturer’s instructions. For total proteome analysis, 100 µg of Fe-NTA flow-through were used. Eluates and flow-through (total proteome) were freeze-dried, solubilized, pooled, and subjected to high-pH reversed-phase fractionation (Thermo Scientific, 84868) before being dried again and reconstituted in 0.1% TFA for LC-MS/MS analysis.

### LC-MS/MS analysis

Both total proteome and phosphoproteome samples were analysed on an Orbitrap Eclipse Tribrid mass spectrometer. Total proteome analyses utilised data-dependent acquisition mode with 180-minute HCD MS2 and 180-minute real-time search (RTS) MS3 methods [78], while phosphoproteome analyses employed 180-minute HCD MS2 and 180-minute MSA SPS MS3 methods as outlined by [79].

### Data processing

Raw mass spectrometry data from both MS2 and MS3 runs were co-processed using MaxQuant versions 2.3.1.0 for TS and 2.5.0.0 for SS against the UniProt human reference proteome database (UP000005640; September 2020 for SS and March 2023 for TS). TMT channel intensities were corrected using batch-specific factors provided by the reagent manufacturer. Searches included variable modification Phospho(STY), with a false discovery rate (FDR) less than 0.01 and a minimum peptide length of six amino acids.

### Data analysis

Processed data were analysed in R using scripts based on ProteoViz [80] and Proteus [81] packages. ProteinGroups.txt and Phospho(STY)Sites.txt files were filtered out for reverse hits, potential contaminants, and proteins identified only by site. Corrected reporter intensities from TMT labelling were normalized using CONSTANd [82] before being log2-transformed for statistical analysis via Linear Models for Microarray Data (limma). Significant changes in total protein and phosphorylation levels were selected using a p value < 0.005.

For sequence motif enrichment analysis the significantly increased phosphosites (p<0.005 and log2FC ≥1) in SEGA samples versus PM or TLE in both trypsin and chymotrypsin datasets were selected and combined. The “Flanking” column was used and duplicated sequences were removed. The amino acid sequences were trimmed down to 11 residues (5 on each side of the phospho-acceptor) and the enrichment motif was plotted using the R package “ggseqlogo”.

### Data availability

The mass spectrometry proteomics data have been deposited to the ProteomeXchange Consortium via the PRIDE [83] partner repository with the dataset identifier PXD069405.

Reviewer access details:

Log in to the PRIDE website (https://www.ebi.ac.uk/pride/) using the following details:

Project accession: PXD069405

Token: EbK9O9NR22FH

### GO analysis

GO analysis was performed using the STRING database V12.0 [51]. All detected proteins or proteins represented by phosphopeptides in the tuber analysis or SEGA analysis were used as the statistical background.

### DNA extraction and genetic analysis

Total DNA was extracted from fresh frozen tuber and SEGA tissues, using the QIAmp DNA Mini Kit (Qiagen, Valencia, CA), following the manufacturer’s instructions. DNA quantification was performed by QUBIT.

The genetic analysis was performed using either one or both of the two approaches (i) a targeted hybrid-capture Massively Parallel Sequencing (MPS) of the entire extent of the *TSC1* and *TSC2* genes, as described previously [84, 85] or (ii) deep sequencing of *TSC2* using multiplex high-sensitivity PCR assay (MHPA) covering 75% of known small pathogenic variants in *TSC2*, including all variant hot-spots, as described previously [86].

The sequencing results were analyzed using our homemade computational pipelines, enabling analysis of single nucleotide variants, indels and large (multiexonic) deletions/duplications, as well as LOH events, as described previously [84, 86].

All identified variants were examined closely in the Integrative Genomic Viewer (IGV). The pathogenicity of the variants was assessed in the TSC LOVD [87] and GnomAD [88] databases.

The HGVS nomenclature was defined using Mutalyzer [89].

### Splicing analyses

#### Data source and access

Paired-end sequencing data were obtained from “The coding and non-coding transcriptional landscape of subependymal giant cell astrocytomas” [56]. Data were accessed from the European Genome-phenome Archive (EGA) under dataset accession EGAD00001005932, via the Data Access Committee EGAC00001001467. The dataset comprises RNA-seq and small RNA-seq from subependymal giant cell astrocytomas (SEGAs; n = 19) and periventricular control brain tissue (n = 8) generated on the Illumina HiSeq 2500 platform.

#### Read processing and quantification

Reads were processed with nf-core/rnasplice v1.0.4 using --aligner star_salmon and --pseudo_aligner false, with GRCh38 primary assembly FASTA and Ensembl GRCh38 release 114 GTF references. STAR and Salmon indices pre-built against these references were supplied to the pipeline. Built-in QC (FastQC, MultiQC) was reviewed prior to downstream analyses.

A dedicated rnasplice run produced exon-level differential usage using DEXSeq with: --dexseq_exon true, --save_dexseq_annotation true. Outputs included exon-bin statistics and per-gene aggregated q-values used for DEU calling at FDR (padj) < 0.05.

Event-type analyses were generated in a separate rnasplice run with rMATS enabled: --rmats true, --rmats_paired_stats false, --rmats_novel_splice_site true, --rmats_min_intron_len 50, --rmats_max_exon_len 1000, --rmats_splice_diff_cutoff 0.05. Significant events were defined at FDR < 0.05, with visualisation focused on events meeting ΔPSI > 0.10.

#### Downstream analyses in R (v4.5.0)

##### DEXSeq (exon-level)

Per-exon DEXSeq results and per-gene aggregated q-values were imported into R. Genes with padj < 0.05 were considered DEU-positive. Volcano plots highlighted exon bins with log2 fold-change > 0.5 and padj < 0.05.

##### rMATS (event-based)

rMATS files were parsed into a unified table for all five event classes. Unless otherwise stated, events with FDR < 0.05 and ΔPSI > 0.10 were considered significant for visualisation.

#### Isoform-level analysis with IsoformSwitchAnalyzeR

Transcript-level quantifications from Salmon were imported via importIsoformExpression, and a combined object was created using importRdata with ignoreAfterPeriod=TRUE (to strip Ensembl version suffixes) and addAnnotatedORFs=TRUE. Low-expression pre-filtering used geneExpressionCutoff=1 TPM and isoformExpressionCutoff=0. Differential isoform usage was tested with isoformSwitchTestDEXSeq. Volcano plots emphasised isoforms with q < 0.05 and ΔIF > 0.10. Where relevant, predicted functional consequences were summarised using analyzeSwitchConsequences(dIFcutoff=0.1) (e.g., NMD status, ORF/UTR changes, domain gain/loss, signal peptide status).

#### Cross-method integration

To compare outputs at the gene level, identifiers were standardised to Ensembl gene IDs. We then defined overlaps across DEXSeq (DEU), rMATS (events), and IsoformSwitchAnalyzeR (isoform switching) and labelled genes supported by ≥ 2 methods as “validated hits.”

#### Gene ontology (GO) enrichment

Over-representation analysis (ORA) for GO Biological Process (BP) used clusterProfiler with Entrez IDs. Background comprised all testable genes detected across methods with results reported as BH-adjusted P values.

#### Computational environment

Nextflow pipelines were executed on King’s College London’s Computational Research, Engineering and Technology Environment (CREATE) high-performance computing (HPC) cluster [90].

### Statistical analysis

Data were expressed as mean +- S.E.M. All data were analysed using Prism 9 (GraphPad). A hypergeometric test was used for analysis of SEGA phosphoprotein overlap with NDD proteins. A one-way ANOVA with Tukey’s post-hoc test was used for multiple comparisons of continuous data P<0.05 considered statistically significant. * *p*≤0.05, \*\**p*≤0.01, *** *p*≤0.001, **** *p*≤0.0001, n.s. non-significant.

## Supporting information

Supplementary figures S1-S3; supplementary tables list

Tables 1-6, spreadsheet format

## Acknowledgements

We are grateful to the TSC Alliance Biosample Repository, the NIH Neurobiobank, King’s College London Neurodegenerative Diseases Brain Bank and the individuals who donated the tissue used in this project.

## Author contributions

Conceptualisation: SKU, DJK, JMB

Proteomics: HF, MS, SRM

Tissue acquisition: LMR, DJK, JMB

Genetic analysis: KK, EBF, DJK

Tissue validation: EA

Computational analysis: MW

Data analysis: MG, KK, JMB

Writing: JMB

Review and editing: All authors

Supervision: SU, DJK, JMB

Funding acquisition: SU, KK, DJK, JMB

