## Supplementary figures S1-S3; supplementary tables list for "The phosphoproteomic landscape of the neurological manifestations in tuberous sclerosis complex"

**Supplemental Figures**

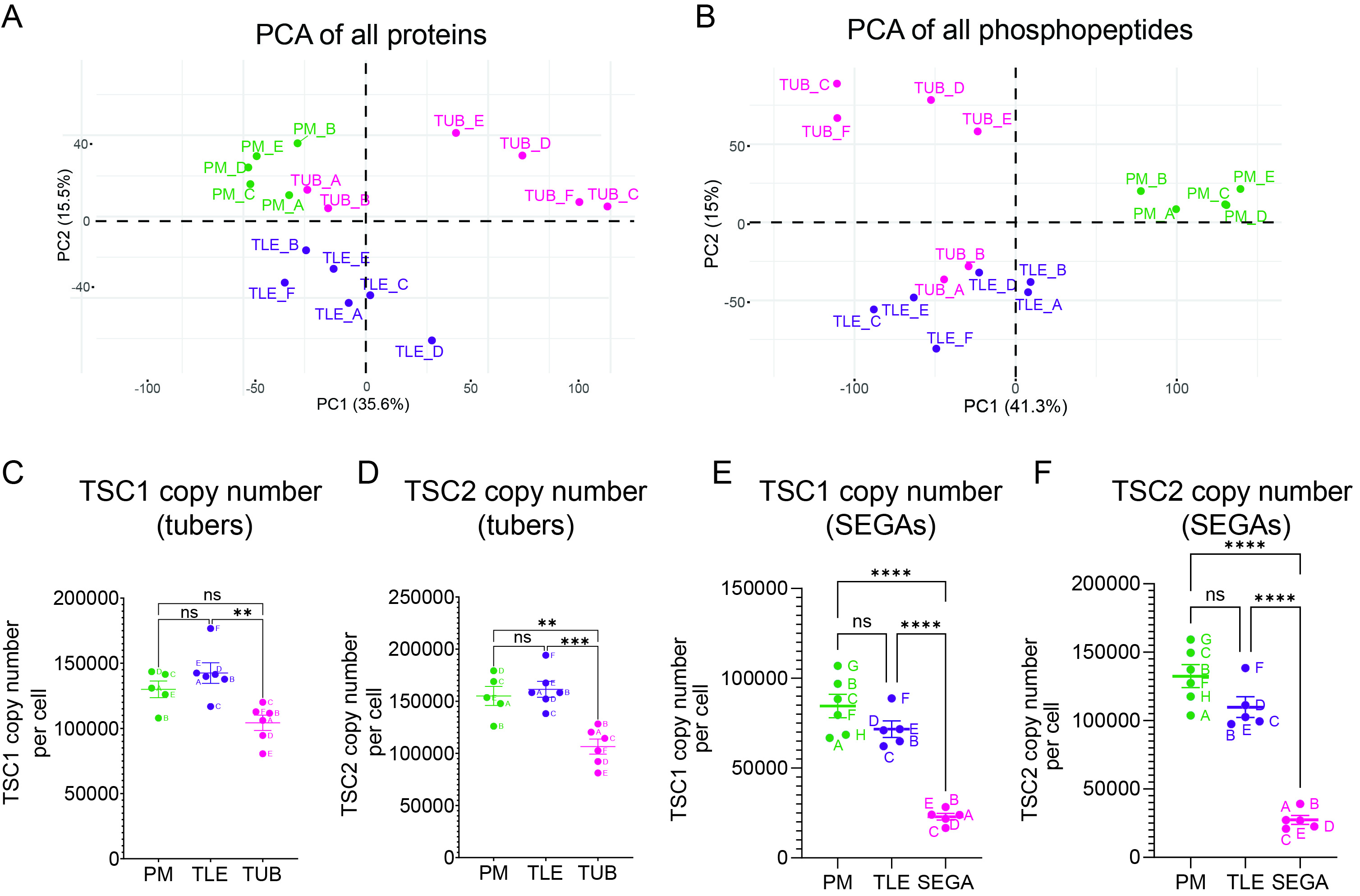

*Figure S1. Quantification of TSC1 and TSC2 protein copy numbers per cell.* (A) PCA plot of postmortem (PM), temporal lobe epilepsy (TLE) and tuber (TUB) proteomic data. (B) PCA plot of PM, TLE and TUB phosphoproteomic data. (C, D) Copy numbers of TSC1 (C) and TSC2 (D) proteins in tubers and control tissue. n=5 for PM, n=6 for TLE, n=6 for TUB. (E, F) Copy numbers of TSC1 (E) and TSC2 (F) proteins in SEGAs and control tissue (trypsin). n=6 for PM, n=5 for TLE, n=5 for SEGA. Data are represented as mean ± SEM and were analysed using one-way ANOVA. ns not significant, * p < 0.05, **p < 0.01, ***p < 0.001****p < 0.0001.

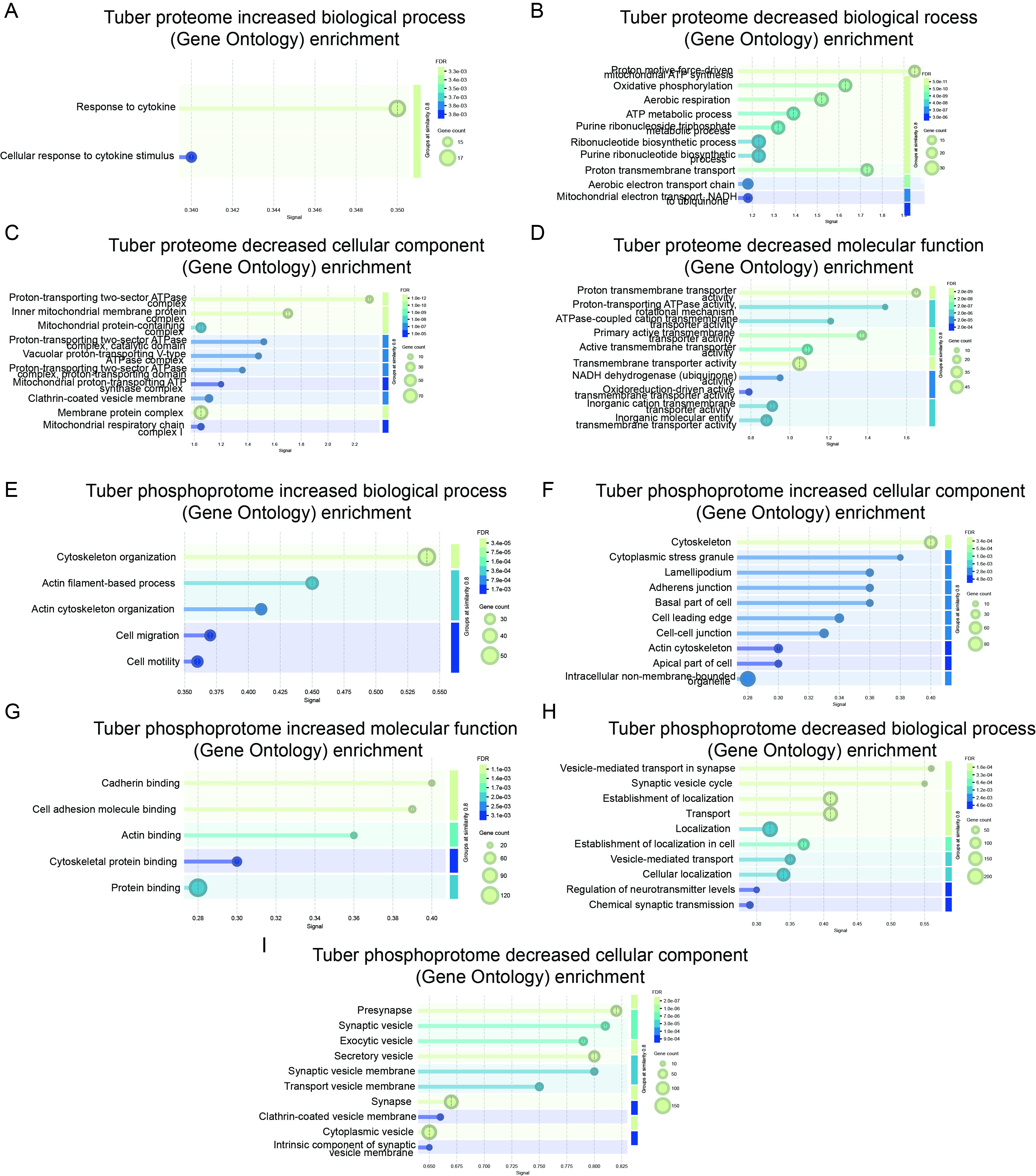

*Figure S2. GO analysis of proteins with altered expression in tubers*. (A) GO analysis of biological process of proteins with increased expression in tubers shows enrichment for cytokines. (B-D) GO analysis of biological process (B), cellular component (C) and molecular function (D) of proteins with decreased expression in tubers shows enrichment for mitochondria-related processes. (E-G) GO analysis of biological process (E), cellular component (F) and molecular function (G) of proteins represented by phosphopeptides with significantly increased expression in tubers shows enrichment for cytoskeleton-related processes. (H, I) GO analysis of biological process (H) and cellular component (I) of proteins represented by phosphopeptides with significantly decreased expression in tubers shows enrichment for synapse-related processes.

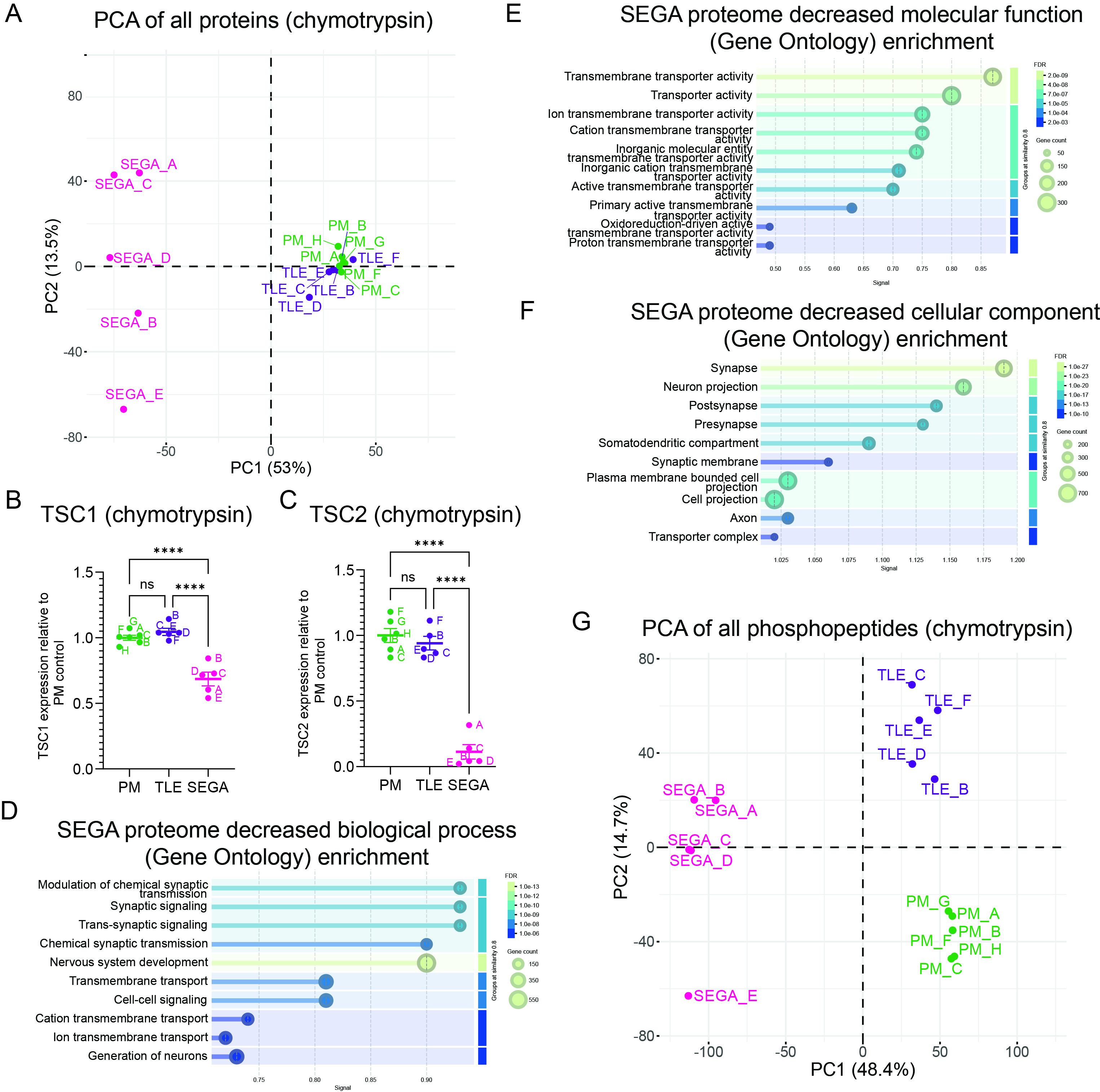

*Figure S3. GO analysis of proteins with decreased expression in SEGAs*. (A) PCA plot of postmortem (PM), temporal lobe epilepsy (TLE) and SEGA proteomic data in the chymotrypsin condition. (B, C) TSC1 (B) and TSC2 (C) protein expression levels are strongly decreased in the chymotrypsin condition. (D-F) GO analysis of biological process (D), molecular function (E) and cellular component (F) of proteins with significantly decreased expression in SEGAs shows enrichment for synapse-related processes. (G) PCA plot of PM, TLE and SEGA phosphoproteomic data in the chymotrypsin condition.

**Supplemental Tables**

*Table S1. Proteomics data for the tuber experiment.*

*Table S2. Phosphoproteomics data for the tuber experiment.*

*Table S3. Direct mTORC1 substrate phosphosites detected in tuber phosphoproteomics.*

*Table S4. Proteomics data for the SEGA experiment (trypsin).*

*Table S5. Proteomics data for the SEGA experiment (chymotrypsin).*

*Table S6. Phosphoproteomics data for the SEGA experiment (trypsin).*

*Table S7. Phosphoproteomics data for the SEGA experiment (chymotrypsin).*

*Table S8. Direct mTORC1 substrate phosphosites detected in SEGA phosphoproteomics.*

*Table S9. SEGA phosphopeptides with significantly increased expression compared to PM or TLE controls (p<0.005, FC ≥ 1) in trypsin and chymotrypsin experiments.*

*Table S10. SEGA proteins represented by phosphopeptides with significantly increased expression compared to PM or TLE controls in trypsin and chymotrypsin experiments.*

*Table S11. Genes with mis-regulated splicing in SEGA tissue identified by at least two analysis pipelines.*

*Table S12. Genes with mis-regulated splicing in SEGA tissue identified by all three analysis pipelines.*
